# Environmentally relevant depleted uranium exposure damages mitochondria, decreases cytosolic reductive capacity, and increases global DNA damage accumulation through a ROS-independent mechanism involving *slingshot protein phosphatase 1b* enrichment

**DOI:** 10.64898/2026.07.02.736169

**Authors:** Phillip H. Kalaniopio, Luke B. Gibbons, Ronald S. Allen, Saoirse M. Matthews, Oscar R. Lujan, Elyès Gaaloul, Cailyn M. Allen, Justin Wilbanks, Connor A. Chassman, Tinna Traustadóttir, Catherine R. Propper, Matthew C. Salanga

**Affiliations:** Department of Biological Sciences, Northern Arizona University, Flagstaff, AZ, USA; Department of Chemistry, Northern Arizona University, Flagstaff, AZ, USA

**Keywords:** metabolism, uranium, arsenic, zebrafish, reactive oxygen species, Nrf2, SSH1

## Abstract

Depleted uranium (DU) is an environmental contaminant with a 30 µg/L (ppb; parts per billion) EPA maximum contaminant level (MCL) for drinking water. The mining of uranium and use of DU in modern weapons underly human exposure that disproportionally impacts military and tribal communities in the United States. Uranium’s radiotoxic characteristics are understood, but its chemical hazards much less so. In zebrafish (*Danio rerio*) and human cell cultures we test the hypothesis that exposure to DU negatively impacts cellular function and development through disruption of mitochondrial metabolism. Using a novel shrapnel model with TEM/SEM+EDS, we showed uranium microparticles caused proximity-dependent mitochondrial disruption. In waterborne exposure paradigms, larval movement was reduced and hatching delayed as a result of reduced movement and not enzyme deficiencies in response to 18 ppb DU, below the MCL. Increased DNA damage accumulation was detected in exposed larva and cells. DNA-damage quantitative PCR of DU-exposed larvae showed increased damage in the *ahr1* locus (nuclear gene) and decreased mitochondrial DNA (mtDNA) copy number, but mtDNA damage levels varied across experiments. Mitochondrial function was assessed using a resazurin-based assay in the presence and absence of antioxidants and showed diminished cytoplasmic reductive capacity. DU exposure alone did not enrich antioxidant gene expression, contrasting with arsenic exposure, a known ROS-inducer and Nrf2-activator. Sulforaphane (SFN), a potent Nrf2-activator, did not blunt the effects of DU exposure, despite activation of antioxidant response element (ARE) genes (*gstp* and *gss)*, but did blunt the effects of arsenic exposure. The most enriched transcript in DU-exposed larvae coded for slingshot protein phosphatase (*ssh*), further exploration revealed *ssh1b* as the zebrafish-specific ortholog activated in response to DU, and inhibition using an identified SSH1 inhibitor, Sennoside A, partially rescued the metabolic and hatching defects observed. Our data points to a cytotoxic mechanism in which DU disrupts mitochondrial function through *ssh1b* enrichment that impairs normal mitophagy, leading to decreased cellular reductive potential independent of either ROS production or ARE-activation. Our results suggest that health impacts from DU exposure may be directly linked to impaired mitochondrial functions.

## 1. INTRODUCTION

Uranium (U) is a naturally occurring and globally distributed heavy metal with a documented hazard profile (Agency for Toxic Substances and Disease Registry (ATSDR), 2013). Uranium mining and the use of uranium-containing munitions have contaminated natural resources, residential areas, and battlefields (Fathi et al., 2013; Faa et al., 2018; Lourenço et al., 2017). In the United States, uranium exposure disproportionately affects two groups: tribal nations and veterans (Papastefanou, 2002; Redvers et al., 2021). While uranium’s radiological hazards are well defined, its role as a chemical toxicant is not fully understood (Ma et al., 2020). Clarifying uranium’s chemotoxic mechanism-of-action is critical for risk assessment and the development of therapeutic or protective approaches, in service to public health.

### 1.1 Mining and Tribal Nations

The Navajo Nation, the largest tribal nation of indigenous Americans, was entangled in the uranium mining vigor of the 1950s, with an estimated 30 million tons of uranium ore extracted from the Nation between 1944 and 1986 (Agency for Toxic Substances and Disease Registry (ATSDR), 2013). Health effects such as increased rates of lung cancer, tuberculosis, and pneumoconiosis were recorded in Navajo miners (Roscoe et al., 1995). The cessation of mining activities was abrupt and did not include engineering controls to “close” mine sites, thus creating a potential environmental contamination source (Ingram et al., 2020). A 2018 study surveyed and quantified uranium content in 82 water sources scattered across the western portion of the Navajo Nation (Jones et al., 2020). Among the 82 water sources sampled, nine exceeded the MCL with a maximum measurement of 560.2 ppb (µg/L) in groundwater (Ingram et al., 2020). Ingestion of water at these levels can lead to deposition in both the kidneys and bone leading to kidney damage by way of acute tubular necrosis, bone damage, reproductive and developmental effects, and the possibility for weak carcinogenicity (Faqir et al., 2025).

### 1.2 Depleted Uranium Munitions

Modern militaries have recognized the utility of depleted uranium (DU) in armor and armor-piercing (AP) munitions, that have contaminated most modern battlefields (Fathi et al., 2013). DU projectiles exhibit a self-sharpening pyrophoric reaction when penetrating a hardened target, that releases uranium oxide microparticles and larger DU fragments into the immediate surroundings (Walkingstick et al., 2018). The microparticles can settle onto the ground where they may enter water resources, contaminate food supplies, or be inhaled when surface soils are disturbed by natural or anthropogenic means (Agency for Toxic Substances and Disease Registry (ATSDR), 2013; Anke et al., 2009). Larger DU fragments in the form of shrapnel may penetrate human tissue that can remain *in situ* for a lifetime with modest mobilization determined by consistent uranium-detection in subjects’ urine (McDiarmid et al., 2025). This exposure mechanism causes tissue proximal to implanted fragments to be subject to DU toxicity as well as supplying a continuous chronic low dose of uranium to other areas of the body and organ systems, such as the skeletal and urinary tract systems (McDiarmid et al., 2021). In a murine shrapnel model, U was measured in the urine, and significant concentrations found in tissue including muscle, spleen, liver, heart, lung, brain, lymph nodes, and testicles, demonstrating that although a shrapnel wound seems to be localized and dormant, DU can be transported and migrate throughout different discrete tissue types (Hahn et al., 2002; Leggett and Pellmar, 2003; Pellmar, 1999).

### 1.3 Depleted Uranium is a Mitotoxin

Mitochondria have emerged as a target of DU chemotoxicity (Ma et al., 2020; Armant et al. 2019). DU exposure disrupts mitochondrial electron transport chain (ETC) complexes, particularly complex IV (cytochrome C oxidase) and complex V (ATP synthase) (Yu et al., 2021). Functional ETC complexes are required to pump protons into the intermembrane space to maintain mitochondrial membrane potential and power oxidative phosphorylation (OXPHOS). DU is linked with the loss of mitochondrial membrane potential (Pourahmad et al., 2006) and decreased ATP synthase activity (Lerebours et al., 2010). Mitochondrial ultrastructure also appears impacted by exposure to DU (Armant et al., 2017). The induction of ROS following DU exposure have been demonstrated *in vitro* and *in vivo* (Gagnaire et al., 2014; Orona and Tasat, 2012; Periyakaruppan et al., 2007), including one study describing a dose-dependent relationship between increased ROS generation and DU (Tasat et al., 2007). Zebrafish exposed to DU exhibit increased ROS production and markers of oxidative stress (Augustine et al., 2012; Gagnaire et al., 2013); however, the molecular interactions by which DU initiates ROS generation remain unclear. In the present study, we test two mutually exclusive hypotheses: one, that DU causes ROS generation independent of mitochondria (*i.e.,* in the cytosol) that impacts the ETC, leading to more ROS production. Two, DU damages mitochondrial structures, leading to ROS production from disrupted mitochondria; that is, ROS production as a symptom of mitochondrial disruption.

### 1.4 Mitochondria and Reactive Oxygen Species

Mitochondria play a crucial role in redox biology for homeostasis and signaling. Mitochondrial dysfunction, such as disrupted ETC activity, diminishment of mitochondrial intermediate molecules or products, changes in mitochondrial ultrastructure, or other mitopathies may impair function (Meyer et al., 2018). OXPHOS constantly produces ROS as a byproduct while electrons are shuttled down the ETC, some leak from ETC complexes or electron carrying molecules (Turrens, 2003). Leaked electrons react with elemental oxygen lacking proper enzymatic guidance, forming superoxide (O_2_^-^) instead of water (Jastroch et al., 2010). Under normal conditions, these species are scavenged by mitochondrial antioxidant enzymes, mainly manganese superoxide dismutase (Mn-SOD; SOD2), and are not pathological (Hong et al., 2024). Disruption of ETC complexes increases both electron leakage rates and superoxide production (Hanukoglu et al., 1993). If the disruption is great enough, a situation may arise in which the rate of electron entry exceeds electron exit and the ETC stalls, causing electrons to accumulate in the mitochondrion, fueling the production of superoxide, and overwhelming mitochondrial antioxidant activities (Zhao et al., 2019). For these reasons, determining whether ROS is a driver or a symptom of mitochondrial disruption is a challenge (Liemburg-Apers et al., 2015).

### 1.5 Slingshot Homolog and Mitophagy

Another mechanism through which DU may be impacting mitochondrial function is through the SSH family of proteins. The family of slingshot homolog proteins are dual specificity phosphatases encoded by *SSH1*, *SSH2*, and *SSH3* in humans, and *ssh1a*, *ssh1b*, *ssh2a*, and *ssh2b* in zebrafish. SSH1, the most widely studied isoform, impairs mitochondrial health and respiration through cofilin-dephosphorylation related to the N-terminal region and impairs mitophagy through a SQSTM1/p62-binding C-terminal region (Cazzaro et al., 2023). Inactivation of SQSTM1/p62 by SSH1 results in accumulation of damaged and ubiquitinated mitochondria, resulting in impaired mitophagy and autophagy (Fang et al., 2021). Basal SSH1 is inactivated due to association with 14-3-3, but oxidation of 14-3-3 causes SSH1 to be released and activated (Bernstein and Bamburg, 2010; Eiseler et al., 2009; Kim et al., 2009). Increased transcription of SSH1 reflected protein expression, was detected in colorectal cancer samples, has been linked to poorer colorectal tumor prognosis in humans, and a SSH1 knockdown model revealed that decreasing expression of SSH1 resulted in attenuation of tumor burden (Song et al., 2020). In zebrafish, SSH1 interacts with VCL and CFL in cardiac myocyte development (Fukuda et al., 2019).

In the present study, we explore the capacity for DU to impair mitochondrial metabolism via either 1) direct ROS generation, or 2) ROS produced *from* mitochondrial dysfunction, as we deduce the mechanistic underpinnings of DU toxicity. In either case, mitochondria-disrupting ROS will oxidatively stress ETC proteins, mitochondrial membranes, and mtDNA, resulting in mitochondrial dysfunction that causes more ROS generation, in an apparent feedback loop (Cadenas and Davies, 2000). Alternatively, mitotoxins can induce mitochondrial dysfunction by mechanisms that do not generate mitochondrial-disrupting ROS (ROS originating chemically in the cytoplasm), and in those instances ROS are predominantly generated from disruption (Hu et al., 2020). In simpler terms, the former puts ROS as both the cause and therapeutic target of U, and in the latter, ROS is a symptom that contributes to other cellular pathologies but is not a tractable therapeutic target for U toxicity.

## 2. METHODS

### 2.1 Zebrafish husbandry and embryo collection

All zebrafish care and use protocols were approved by Northern Arizona University’s Institutional Animal Care and Use Committee (IACUC), protocol number 23-009. Zebrafish were maintained in Tecniplast ZebTec Active Blue^TM^ systems. Semi-recirculated system water was continuously monitored while recirculated to control water temperature (28.5 °C ± 0.5), pH (7.5 ± 0.25), and conductivity (700 ± 100 µSiemens). The systems are maintained in a 14/10-hour light-dark cycle, and fish were fed twice daily with Skretting Gemma® 75, 150, or 500, depending on size, and supplemented 3-5 per week with a diet of live rotifers (*Brachionus plicatilis*) or brine shrimp nauplii (*Artemia salina)* depending on size (Lawrence et al., 2016). Zebrafish embryos were obtained by spawning adult zebrafish (∼5-18 mpf) in breeding tanks with perforated inserts that allow embryos to pass through but not adults. Embryos were collected from the breeder tanks and rinsed with 0.3x Danieau’s solution (DanS) and disinfected using 0.01% Clorox bleach 1 min soak then 5 X 5 min washes in 0.3x DanS. Surface disinfected embryos are then moved to 15 cm petri dishes and maintained in an incubator set to 28.5 °C and a 14/10-hour light/dark cycle.

### 2.2 Chemical preparation

Depleted uranium in the form of triuranium octoxide powder (U_3_O_8_; CAS#1344-59-8) was gifted to us by Dr. Jani Ingram (Regents’ Professor, Northern Arizona University), and originally procured from Strem Chemicals (Newburyport, MA). For implantation experiments, U_3_O_8_ was made into a supersaturated slurry with ultrapure water and stored in amber 1.5 mL Eppendorf tubes. For waterborne exposure paradigms, depleted uranium in the form of uranyl nitrate (UN; (UO_2_(NO_3_)_2_) 6·H_2_O; CAS# 13520-83-7, Electron Microscopy Sciences, CAT# 22600), a soluble uranium species, was used to prepare solutions of 0.3, 3, 30, 49.7, and 300 µg/L UN or 0.18, 1.81, 18.1, 30, and 181 ppb uranium atoms in 0.3x DanS; early dose-finding experiments feature a dose curve consisting of 0, 23.8, 238, or 2,380 ppb U (0, 0.1, 1, 10 µM UN), prepared in the same manner. Sodium (meta) arsenite (NaAsO₂) was obtained from Sigma Aldrich (CAT# S7400; CAS# 7784-46-5, Sigma Aldrich), and a 1 mM stock solution was prepared by dissolving the arsenic powder in ultrapure water. Next, a working solution was prepared by diluting the 1 mM stock solution with water and 30x DanS to a final concentration of 1 µM NaAsO₂, in 0.3x DanS (approximately 37.5 ppb arsenic). Zebrafish SFN exposure studies were conducted using DL-sulforaphane N-acetyl-L-cysteine (Cayman Chemicals, CAT# 16098, CAS# 334829-66-2), which was reconstituted in 100% DMSO to a stock concentration of 40 mM, aliquoted, and frozen at -20 °C until use. MitoTEMPO was acquired from Sigma-Aldrich (CAS#1334850-99-5), dissolved in ultrapure water to a stock concentration of 20 mM, aliquoted, and frozen at -20 °C until use. TEMPOL (4-hydroxy-TEMPO) was acquired from Sigma-Aldrich (CAS# 2226-96-2) and dissolved in 100% DMSO to a stock concentration of 100 mM, aliquoted, and frozen at -20 °C until use. Dithiothreitol (DTT; CAS# 3483-12-3) was acquired from Fisher Bioreagents and dissolved in water or 0.3x DanS. Sennoside A (CAS# 81-27-6) was obtained from Cayman Chemicals (Ann Arbor, MI, USA) and diluted to a stock concentration of 1 mM in 1x phosphate-buffered saline (PBS). This stock was divided into 1.5 mL amber Eppendorf tubes and froze at -20 °C until use. Brusatol (CAT# 308783, Cayman Chemical) was obtained and immediately diluted into 100% DMSO to produce 1 mM stock solutions in amber 1.5 mL Eppendorf tubes, frozen at -20 °C until use.

### 2.3 Microsurgical Implantation of DU

We devised a novel technique to emulate chronic DU exposure by shrapnel wounds in zebrafish. Embryos were collected from AB strain wildtype (WT) fish crossed with triple mutant Tg:mitf:BRAF^V600E^; *mitfa^-/-^*; *p53^-/-^* producing triple heterozygote offspring. Initially, we wanted to know if the micro-implanted U_3_O_8_ increased cellular neoplasia; thus, our selection of the triple heterozygotes with oncogenic potential. Our question changed, as DU’s impact on mitochondria emerged as our primary question (Allen Jr., 2021). Thus, it was determined that heterozygosity was not relevant for our analysis. At 3-4 dpf, embryos were randomly divided into equal-sized groups, manually dechorionated, and divided into two groups receiving either a depleted U_3_O_8_ implant or sham surgery. Embryos were anesthetized with 0.0016% w/v tricaine (Ethyl 3-aminobenzoate methanesulfonate; CAS# 886-86-2, buffered with Na_2_HCO_3_ to pH 7.2) and placed on prepared surgical plates lined with 2% agarose gel, infused with 0.4% methylene blue, and submerged under a thin layer of 0.3x DanS with their left flanks nearest to the surface (Fig. 2a). An electrolytically sharpened tungsten-wire needle was used to implant DU into the left tail flank, posterior and dorsal to the tip of the yolk sack extension – a process akin to tattooing (Fig. 2b). Immediately following implant, individuals were evaluated by monitoring heart activity and blood flow of major vessels using light microscopy (Fig. 2c). Embryos were then moved to new petri dishes with fresh 0.3x DanS overnight and nurtured under normal zebrafish housing conditions until sampling at approximately 28 dpf.

### 2.4 Waterborne treatment*s*

Waterborne exposures were performed in 10 cm petri dishes, and treatment compounds were diluted to working strengths in a 0.3x DanS solution supplemented with vehicle controls when necessary.

#### 2.4.1 Uranyl nitrate and sodium meta-arsenite

Exposure solutions were made from 10x concentrated stocks (as described in **2.2)**, diluted in 0.3x DanS to the appropriate treatment concentrations, and refreshed by exchanging ∼50% volume every day of exposure.

#### 2.4.2 Sulforaphane

Larval zebrafish were plated in 0.3x DanS with increasing volume of 40 mM SFN stock solution in 100% DMSO in petri dishes to create 1, 5, 10, and 40 μM SFN final concentrations. DMSO was added to treatment groups and controls to maintain a final concentration of 0.1% v/v DMSO for all groups and treatment solutions were refreshed daily. Zebrafish subject to SFN pretreatment were plated in SFN-containing media within 2 hpf. During the 2 hpf to 1 dpf timeframe, zebrafish embryos were exposed to DU-free control or SFN-containing DanS. At 1 dpf, media change for metal-containing treatment groups included incorporating DU or As into exposure conditions. SFN groups were exposed to SFN from 2 hpf through 3 dpf.

#### 2.4.3 TEMPOL

Larval zebrafish were exposed in a similar manner to sulforaphane (**2.4.2**), as TEMPOL is soluble in DMSO, but they were not subjected to a one-day drug-only pretreatment regime as in the case of SFN. 100 μM was chosen based on previous efficacy of this dose in zebrafish (Guan et al., 2021). TEMPOL was administered at a final concentration of 100 μM and final volume of 25 mL to petri dishes containing DanS and relevant concentration of DU once daily, from 1 to 5 dpf.

#### 2.4.4 MitoTEMPO

MitoTEMPO is water-soluble, leading to the elimination of DMSO-vehicle control groups for comparison of this antioxidant. MitoTEMPO was administered at a final concentration of 40 μM and final volume of 25 mL to petri dishes containing DanS and relevant concentration of uranium once daily, from 1 to 5 dpf.

#### 2.4.5 Sennoside A

Final treatment solution preparation and exposure consisted of dilution of the 1 mM stock to relevant treatment concentration in 0.3x DanS and was refreshed once daily until experiment completion.

#### 2.4.6 Brusatol

1mM Brusatol stock was diluted to a final concentration of 500 nM, based on previous efficacy (Bian et al., 2023; He et al., 2023; Zhang et al., 2024), in 0.3x DanS and refreshed once daily until experiment completion.

### 2.5 Cathepsin L ELISA

The SensoLyte Rh110 Cathepsin L Fluorometric Assay Kit (AnaSpec; CAT# AS-72217) was used to quantify cathepsin L enzyme activity in zebrafish embryos. Briefly, cathepsin L enzyme cleaves a non-fluorescent substrate producing brightly fluorescing rhodamine 110 that can be measured using a microplate reader. A cathepsin L inhibitor (component E) included with the kit was used as a negative control. Zebrafish embryos were collected between 2-3 hpf and exposed to 0, 23.8, 238, or 2,380 ppb U (0, 0.1, 1, 10 µM UN) until 48 hpf, at which time UN was removed, embryos were washed twice with 0.3x DanS before transferring to individual microcentrifuge tubes containing 100 µL of ice-cold 1x PBS. Next, the samples were homogenized by sonication (15 sec/20% amplitude; Fisher Scientific, Model 120 Sonic Dismembrator). A 96-well black-walled clear-bottom plate was prepared with each well containing either 100 µL of Milli-Q water, or 50 µL of sample + 50 µL of fluorescent substrate (across five replicates), or 50 µL of sample homogenate + 50 µL fluorescent substrate + 10 µL of Component E (Cathepsin L inhibitor). The plate was incubated at 28.5 °C for one hour and then read at 485 nm excitation/525 nm emission using a microplate reader (BioTek; Synergy HTX).

### 2.6 Fish Embryo Toxicity Test (FET)

Embryos (approx. 64-cell stage) were incubated in glass petri dishes containing treatment solutions. Twenty-four hours later, the embryos were inspected and individually transferred to a 24-well plate (CLS3738, Millipore Sigma), with each well containing 2 mL of the same treatment solutions. Embryos were assessed by stereomicroscopy every 24 hours for 96 hours total with a series of endpoints scored to determine lethality or developmental impairment: *e.g.,* detachment of tail, somite formation, eye formation, movement, blood circulation, heartbeat, pigmentation of head and body, pigmentation of the tail, yolk extension nearly empty, pectoral fin, protruding mouth, and hatching (OECD, 2013, Beekhuijzen et al., 2015). Teratogenic endpoints include malformations of the head, sacculi/otoliths, tail, heart, body shape, and yolk sac (Beekhuijzen et al., 2015). Using a 200 μL pipette tip (USA Scientific, Florida), embryos were given up to three light touches to see if they responded with movement and to adjust their position for imaging. Plates with embryos were incubated at 28.5 °C with a 14-light and 10-dark hour lighting system (3915LT, Thermo Fisher Scientific, Massachusetts). At 96 hpf, a final assessment was made, and all remaining larvae were euthanized. Endpoints were scored based on specific developmental criteria and point systems (OECD, 2013).

### 2.7 Movement Observations

A DanioVision (Noldus; Netherlands) observation system was used to track and record intra-chorionic movement of exposed zebrafish. Embryos were collected at roughly 3 hpf and evenly dispensed into petri dishes containing 0.3x DanS supplemented with treatment solution. Embryos were raised in these common gardens for 24 hours, then healthy zebrafish were individually moved to 24-well plates and maintained at 28.5 °C. At 48 hpf, the plate was mounted on the DanioVision stage with the chamber recapitulating normal housing parameters (*i.e.,* 28.5 °C and 14/10 light/dark). Test subjects were recorded for 22 hours, and the recordings were analyzed for the total distance moved before hatching by marking the center of the chorion in the image with a red pinpoint; the movement of the red pinpoint was then used to compute distance moved.

### 2.8 Citrate Synthase Specific Activity

At 3 dpf, embryos exposed to DU (n = 6 groups of 15 – 30 fish per group) were homogenized in 20 µL ice-cold enzyme extraction buffer (20 mM HEPES pH 7.17, 1 mM EDTA, 0.1% Triton X-100) using a handheld homogenizer (CAT# 15-340-167, FisherBrand). Homogenates were centrifuged at 4,000 x g for 15 min at 4 °C, and the supernatant was transferred to new tubes and kept on ice for the duration of the experiment. The assay was conducted following published protocols (Janssen and Boyle 2019). Briefly, in the presence of citrate, 5,5’-dithiobis (2-nitrobenzoic acid) (DTNB) is converted to 5-thio-2-nitrobenzoic (TNB), which is detectable by measuring the absorbance at 412 nm. Protein was quantified using the RC/DC assay (BioRad) and normalized amounts of protein extracts were added to a reaction buffer containing 50 mM KPI buffer (pH 7.4), 100 µM DTNB (D8130, Sigma-Alrdich), and 115 µM acetyl CoA (A2181, Sigma-Aldrich). Measurements were performed in a clear-bottom, dark-walled 96-well microplate (CAT# 07-200-565, Fisher Scientific) with a final reaction volume of 200 µL. Absorbance at 412 nm and 29 °C was recorded for 3 min at 30 sec intervals using a BioTek Synergy H1 plate reader (Agilent) to temporally measure reduction of DTNB by citrate synthase (CS), followed by addition of 5 mM oxaloacetate to a final concentration of 100 µM and a subsequent run with the same specifications as the previous read. CS specific activity was normalized to protein content and expressed as nmol/min/mg protein as follows:

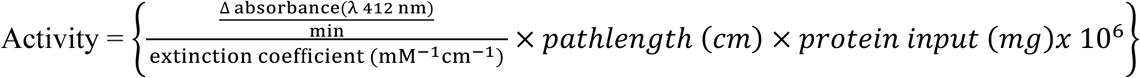

### 2.9 Transmission Electron Microscopy

Transmission electron microscopy (TEM) was used to investigate ultrastructural changes in cells near or far from DU implants. Fish from U_3_O_8_ and sham surgery cohorts were euthanized and immediately fixed in 2.5% glutaraldehyde (CAT# 16200, Electron Microscopy Sciences) with sodium cacodylate trihydrate buffer (over three nights at 4 °C with mild agitation on a rocker or rotator (Fig. 2). Samples were rinsed at RT (∼21 °C) with Na-cacodylate three times for 15 min each. The samples were then fixed with 1% osmium tetroxide for one hour (CAT# 19100, Electron Microscopy Sciences). Samples were then washed with distilled water three times for 15 min each. A graded EtOH dehydration was performed with 1×30%, 1×50%, 1×70%, 1×95%, and 3×100% EtOH for 20 min per step. Spurr’s resin (CAT# 14300, Electron Microscopy Sciences) was used to perform a graded infiltration using 1:3, 1:1, and 3:1 Spurr’s:EtOH, followed by 3×100% Spurr’s for 1 hour per each infiltration step, followed by embedding in a flat mold (Ted Pella Inc., CAT# NC0561014), and curing at 60 °C for one week. Sample blocks were thick-sectioned (10 μm) with a DiATOME® Histo 8 mm 45° knife (CAT# 123-DK4-H80) until a putative DU implant could be visualized by light microscopy and staining with toluidine blue (CAT# DcU-10, Electron Microscopy Sciences, CAS # 92-31-9), then ultrathin sectioned (0.06 μm) using a Leica EM UC6 ultramicrotome and a DiATOME® Ultra 3 mm 45° knife (CAT# 123-UDK4-W30). Floating ultrathin sections were collected onto formvar-coated slot grids and stained with UranyLess EM Stain (CAT#s 2010-Cu and 22409, respectively; Electron Microscopy Sciences) for 20 min, rinsed in distilled water four times, then lead citrate for 10 min, rinsed in CO_2_-free ethanol twice and CO_2_-free water twice. Stained sections were viewed using a JEOL JEM-1200EX-II TEM with integrated digital image capture.

### 2.10 Image scoring

Raw *.tif files of electron micrographs were blinded and assigned randomly generated numerical identifiers via Microsoft Excel. A key was kept separate from any evaluator’s access. Mitochondria in each image were cropped individually to remove any positively identifying source surrounding the organelle (*e.g.,* visible DU deposits, landmarks, etc.). Each image of a mitochondrion was then copied three times, and unique numerical identifiers were assigned using the random number generator (Fig. 2e). All images were then shuffled and made into a catalog. Examples of healthy vs. unhealthy mitochondria from Bourrachot et al. (2014) were placed at the front of the catalogue and evaluators were instructed to consider those first images as reference points. Four evaluators were presented with the catalogue individually on an electronic screen and asked to score mitochondrial health on a scale of 1-5 (1 unhealthy to 5 healthy). Scores for each image were then averaged together, unblinded, and grouped by proximity to DU deposit (Fig. 2e-f). Mitochondria that were ranked within one degree across the three technical replicates were deemed acceptable. In contrast, mitochondria showing more than one rank difference across the three replicates were deemed too variable and not included in the summary statistics.

### 2.11 Energy-Dispersive X-ray Spectroscopy (EDS)

EDS was used to confirm the presence of DU in ultrathin sections after TEM imaging. This method was possible thanks to the use of Uranyless heavy metal stain (as opposed to the use of traditional uranyl acetate). Grids were placed on trimmed index cards, situated on aluminum staging studs with minimal overhang and fixed in place using copper tape. The sections were then sputter-coated with gold-palladium alloy for 5 seconds before analysis in a Zeiss Supra 40VP field emission scanning electron microscope with a Thermo Scientific UltraDry EDS. The electron beam (0.5 kV accelerating voltage) was positioned over the section on the grid, and EDS analysis was used to confirm the presence of elemental uranium atoms in sections from sham or implanted fish.

### 2.12 PBMC Isolation

Whole blood was taken via donor antecubital draw into EDTA vacutainers (CAT# 367835, BD), diluted 1:1 with room temperature 1x PBS in a 50 mL conical tube (CAT# 14-432-22, Falcon^TM^), inverted, and loaded into SepMate^TM^-50 (IVD) tubes with Lymphoprep^TM^ (STEMCELL Technologies) density gradient. SepMates were centrifuged for 15 min @ 1200 x g, additional plasma was removed so 2 - 3 mL of plasma was left above the white buffy coat layer, and cells were carefully poured into a 15 mL centrifuge conical tube, being careful not to invert the SepMate for longer than two seconds. The 15 mL tube containing cells was centrifuged for 10 minutes at 400 x g, supernatant discarded, and cells resuspended in 3 mL of 1:1 1x PBS:0.4% trypan blue (CAT# T10282, ThermoFisher). Cells were counted on a Countess 3 Automated Cell Counter (ThermoFisher) using cell counting chamber slides (CAT# C10283, ThermoFisher), diluted to desired concentrations using RPMI 1640 (R8758, Sigma-Aldrich), and used immediately.

### 2.13 HEK 293 Culturing

Human embryonic kidney (HEK 293) cells purchased from ATCC (CRL-1573, ATCC, Manassas, Virginia) were cultured in T-75 tissue culture treated flasks in Dulbecco’s Modified Eagle Medium (1x DMEM, CAT# 21063-029, Gibco) supplemented with 10% Fetal Bovine Serum (FBS; CAT# A56704-02; Gibco) and 1% penicillin/streptomycin (CAT# 30-2300, ATCC) to produce Complete DMEM (CDMEM). The following day, cells were incubated for 24 hours with CDMEM alone or CDMEM conditioned with either 1) DMSO, DU, or As with and without 40 µM sulforaphane or 2) increasing doses of DU.

### 2.14 Resorufin-based fluorescence assays (alamarBlue)

Wildtype (AB, Tü, and TL) zebrafish larvae were used to assess metabolic function in response to DU exposure using alamarBlue adapted from Renquist et al., 2013 (Renquist et al., 2013). Larvae were exposed to DU or control solutions from 2 hpf to 3 dpf. Any unhatched larvae at 4 dpf were manually dechorionated with fine forceps, and larvae were then rinsed three times with 0.3x DanS. Each larva was subsequently transferred to one well of a 96-well, black-walled clear bottom Corning plate (used above in **2.8**). Excess DanS was removed and immediately replaced with 150 μL of alamarBlue reaction solution. The alamarBlue reaction solution was prepared using Invitrogen alamarBlue^TM^ High Sensitivity reagent (CAT# A50101, ThermoFisher), 0.3x DanS and addition of sodium bicarbonate as an additional buffering agent. At time point zero, immediately following the addition of alamarBlue, the fluorescence was measured using a fluorescence plate reader (Agilent Synergy H1 Multimode Plate Reader) at excitation/emission of 530/590 nm. Microplates were covered with aluminum foil to block any light exposure and incubated at 28.5 °C ± 0.5 for 24 hours. Following the incubation period, the plate was read a second time (*i.e.,* timepoint 24-hr).

#### 2.14.1 Dithiothreitol (DTT) alamarBlue validation

Four DTT solutions were made in ultrapure water (34.1, 17.05, 1.705, 0.1705 mg/mL). 20 µL of each DTT dilution solution was added to several wells of a dark-walled 96-well microplate (Corning). Then, 200 µL of alamarBlue reaction solution was added, the plate was incubated at room temperature for 1 hour, and fluorescence emissions were measured with blanks subtracted (Suppl. Fig. 1).

#### 2.14.2 Cell culture alamarBlue

For PBMC alamarBlue, PBMCs were isolated as in **2.11**, counted, and plated at a density of 100,000 cells per well with RPMI 1640 immediately prior to the addition of alamarBlue dye. Fluorescence was subsequently read at λ 570/600 nm at 0 and 2 hours post-addition of dye, with results reported as a percentage normalized to the change in fluorescence of the control group. For HEK 293 alamarBlue, cells were seeded on a 12-well plate to 15% confluence with media (CDMEM), 10 µL of alamarBlue dye was added directly to wells and fluorescence was subsequently read at λ 570/600 nm at 0 and 2 hours post-addition of dye, with results reported as a percentage normalized to the change in fluorescence of the control group.

#### 2.15 Comet assay for DNA damage detection

2.16 Zebrafish larvae were raised to 5 dpf in 0.3x DanS only or exposed to increasing doses of DU. On day five, larvae were euthanized and transferred to 1.5 mL Eppendorf tubes (CAT# 022431021, Eppendorf), and 100 µL of larval dissociation mix composed of 300 µg/mL collagenase (C9891; Sigma Aldrich) and 0.235% Trypsin-EDTA (25200-056, Gibco). Each tube received 100 µL of dissociation mix and was thoroughly mixed by trituration at room temperature < 1 min per tube. Dissociation was then neutralized with the addition of 900 µL DMEM + 10% fetal bovine serum. Cell suspension was filtered through a 70 µm cell strainer (CAT# 352350, Falcon) into a fresh 1.5 mL Eppendorf tube and centrifuged for 30 seconds at 1000 rpm. Pelleted cells were resuspended in 1 mL of DMEM + 10% FBS and immediately put on ice until agarose embedding. Frosted microscope slides are precoated with high-melting point agarose and allowed to solidify at 4 °C. Cell suspensions (in 1x PBS) are mixed with low-melting point agarose (37 °C), dispensed onto precoated slides, and immediately coverslipped. Slides were allowed to harden at room temperature before immersion in 4 °C lysis buffer (4250-050-01, R&D Systems) in the dark for 30–40 minutes. Electrophoresis conditions for alkaline comet: slides are incubated in alkaline buffer (pH > 12; NaOH and EDTA-based) for DNA unwinding prior to electrophoresis. For neutral comet: slides are placed directly into neutral electrophoresis buffer (300 mM sodium acetate, 100 mM Tris-HCl, pH 8.3). Slides are electrophoresed at 20 V (constant voltage) for 25 minutes at 4 °C in the dark. All slides within a single experiment are processed simultaneously to minimize variability. After electrophoresis, slides are washed in milliQ-water and stained with SYBR Gold nucleic acid stain (S33102, Invitrogen) for 10–15 minutes at 4 °C in the dark. Slides are rinsed, coverslipped, and imaged using an epifluorescence microscope (Zeiss AX10 with digital camera and fluorescent filter cube with excitation/emission 450-490 nm/515-565 nm, respectively). Consistent lamp intensity and exposure settings are maintained across experiments to reduce signal variability and fading. All slides from a single experiment are imaged within a 4-hour window. Slides are systematically scanned to capture non-overlapping, in-focus comets across the entire coverslipped area. Images are randomized and blinded prior to scoring using R (R version 4.5.0) or ImageJ-based blinding tools. Comets are quantified using CometScore software, and OTM values are exported for statistical analysis in GraphPad Prism (described below).

### 2.17 Semi-Long Run quantitative PCR for DNA damage detection (SLR qPCR)

Semi-long run (SLR) qPCR was performed as described in (Zhu and Coffman 2017). Briefly, 5 dpf larvae were sacrificed, and total gDNA was extracted and purified using a Monarch Genomic DNA Purification Kit (T3010, New England Biolabs; Ipswich, MA). Genomic DNA concentration and A260:A280 ratios were determined using a Nanodrop 2000 spectrophotometer (ThermoFisher; Waltham, MA). DNA samples with A260:A280 ≥ 1.8 were stored at -20 °C, and samples with A260:A280 < 1.8 were cleaned and concentrated (D4013, D4033, Zymo; Orange, CA) and re-measured. Cleaned and concentrated samples meeting the A260:A280 ≥ 1.8 threshold were retained, and those that did not were discarded. Genomic DNA extracts served as templates for SLR. Real-time SLR qPCR was carried out in a Bio-Rad CFX384 Real-Time PCR Detection System (Bio-Rad, Hercules, CA). Reactions were assembled with 6.4 ng template DNA, 1x Syto-9 (S34854, ThermoFisher; Waltham, MA), 1x dNTPs, 1x Q5 reaction buffer, 1x Hotstart Q5 polymerase (N0447, B9027, M0493, respectively, New England Biolabs; Ipswich, MA), and NF-H_2_O, for a final reaction volume of 10 µL per replicate well. Primer sets and reaction conditions are listed in (Suppl. Table 1). Melt analysis was performed at the end of each run. Primer efficiency was evaluated by absolute quantitation through the application of a standard curve made up of serially diluted templates (1:10-1:1,000,000). No-template controls (NTC), and three to four technical replicates per sample were used for every run. DNA damage was quantified as previously described (Lehle et al., 2014; Zhu and Coffman, 2017).

### 2.18 Total RNA extraction

DNA-free RNA was extracted using a Direct-zol RNA MicroPrep kit (CAT# R2060, Zymo Research) following the manufacturer’s instructions. Briefly, 6-10 larvae per treatment groups were harvested at 3 dpf, euthanized by rapid cooling in <4 °C DanS, and transferred to 1.5 mL Eppendorf tubes containing 500 μL of TRIzol™ reagent (CAT# 15596018, Invitrogen) and 10 - 20 ceramic beads (diameter = 1.4 mm). The samples were agitated at 2,000 RPMs for one hour in an Eppendorf thermomixer set to room temperature. Next, 500 μL of pure ethanol was added to each tube and mixed by gentle inversion. The solution was moved through the binding column per the manufacturer’s instructions. DNA-free RNA was eluted with 10-50 μL NF-H_2_O, into fresh 1.5 mL tubes, and quantified using a NanoDrop® 2000 (ThermoFisher Scientific). RNA was either immediately reverse transcribed into cDNA or stored at -80 °C for later use.

### 2.19 cDNA synthesis

DNA-free RNA template and iScript cDNA Synthesis Kit (CAT# 1708890; Bio-Rad) were used for cDNA synthesis following the manufacturer’s instructions. Reaction mixes were prepared using 0.5-1 μg of total RNA template, and a SimpliAmp™ thermocycler (Applied Biosystems) to control incubation times. Freshly synthesized cDNA was used immediately in PCR or stored at -20 °C for later use.

### 2.20 Quantitative PCR

Real-Time quantitative PCR was performed on a Bio-Rad CFX384 Touch Real-Time PCR detection system with CFX Maestro® software (version 4.1.24331219, Bio-Rad, Hercules, California). 5-10 µL reactions were prepared from fresh master mixes of SsoAdvanced Universal SYBR Green Supermix (CAT#1725271, Bio-Rad) with 10 µM of forward and reverse primers, according to the manufacturer’s instructions. Reactions were loaded into a 384-well hard-shell thin-walled PCR plate (HSP3805, Bio-Rad) and sealed with Microseal ‘C’ PCR Plate sealing film (MSC1001, Bio-Rad). Reactions were run in technical quadruplicate for each biological replicate, using a two-step thermal cycling method. Melt analyses were performed to confirm amplification specificity. Expression levels were calculated using the built-in ΔΔcT analysis Gene Study method, normalized using the geometric mean of the two housekeeping genes, *actb* and *ef1a* (Supp. Table 1). Primer efficiencies were determined by standard curves using cDNA made from age-matched unperturbed larvae, serially diluted 1:5 across five orders of magnitude.

### 2.21 Statistical Analysis

Statistical analysis and graph production were completed in Prism GraphPad (Version 10.4.2). Data was screened for normality, then outlier values were removed using one of the two following methods: (i) Sample values further than two standard deviations from the mean were deemed outliers and excluded from analysis; (ii) Sample values were assessed using the Robust Outlier tool (ROUT; Q = 1.0; Prism GraphPad™ Vers: 10.4.0), and outliers excluded. Parametric data were analyzed by one-way ANOVA and Tukey’s post-hoc test or a Brown-Forsythe ANOVA followed by a Dunnett’s post-test when standard deviations were unequal. Non-parametric data were analyzed by Kruskal-Wallis analysis of variance and Dunn’s posttest for pairwise comparisons. A p-value threshold of 0.05 (⍺ = 0.05) was used to determine significance, with values ≤ 0.05 deemed statistically significant. Movement data (as repeated measures) were analyzed by two-way Mixed Model ANOVA and Dunnett’s test for comparisons.

### 2.22 Elemental Quantification of exposure solutions

50 mL of exposure solution (0.3, 3, 30, 49.7, 300, 3000) with relevant solvent controls (0.3x Danieau’s Solution, MilliQ-water used for exposure solution preparation, and LC-MS grade water as a technical blank) were collected in 50 mL Falcon tubes (CAT# 14-432-22, Falcon^TM^). Sample concentrations of ^238^U were analyzed following EPA Method 6020 using an Agilent 8900 inductively coupled plasma mass spectrometer (ICP-MS). The system was equipped with an SPS-4 autosampler, peristaltic pump sample introduction system, a MicroMist glass concentric nebulizer, x-Lens ion lenses, and an electron multiplier detector (all from Agilent). ^238^U was analyzed in single-quadrupole mode with helium cell gas to resolve spectral interferences. Samples were diluted with 2% nitric acid (CAS# 7697-37-2, HNO_3_, VWR Aristar Ultra) to fit within the calibration range. Method blanks were run alongside samples, and triplicate measurements were taken for all samples, standards, and blanks. A ^103^Rh internal standard solution was introduced on-line through the peristaltic pump system. Internal standard recovery was between 80-120%. External standard calibration solutions, from 0 to 5 ng/L, were prepared using a multi-element standard (Part No. CMS-1-125ML, Inorganic Ventures CMS-1). The calibration curve for ^238^U had an R^2^ value greater than 0.999. A certified reference material (NIST SRM 1640A: Trace Elements in Natural Water) was used as a quality control standard and was measured within the range of the certified concentration value. Final concentrations are displayed in Supplemental Figure 4 relative to their calculated values (Suppl. Fig. 4)

## 3. RESULTS

### 3.1 Depleted uranium delays normal hatching by decreasing intra-chorionic movement

To broadly define the effects of low-level DU exposure on zebrafish development, we performed a set of modified FETs. Zebrafish embryos exposed to 23.8, 238, and 2380 ppb U (1, 10, 100 µM) of uranyl nitrate hatched approximately 24 hours later than untreated sibling controls (Fig. 1A). To determine if this delayed hatching was a consequence of impaired hatching enzyme synthesis, we performed an ELISA to measure cathepsin L activity, a primary hatching enzyme; embryos treated with DU showed normal cathepsin L activity compared to control animals (Fig. 1B). We next determined whether intra-chorionic movement was impaired in DU-exposed embryos using real-time video from 72-96 hpf which includes the normal hatching window for untreated and delayed larvae 72-83 hpf. Intra-chorionic movement significantly decreased in DU-exposed larvae during this period, with all control larvae having hatched by 82 hpf (Fig. 1C).

**FIGURE 1.**
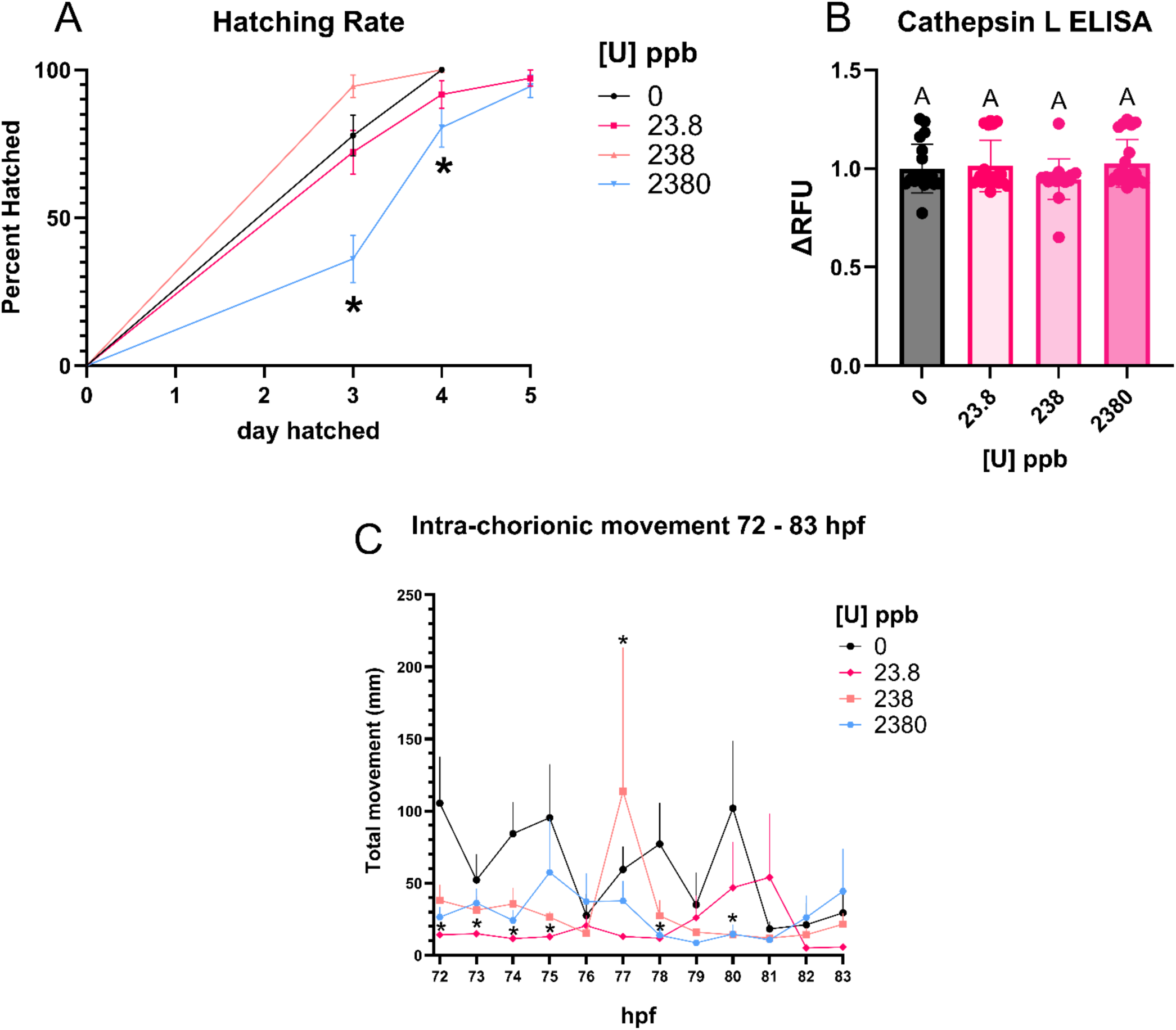
Depleted uranium delays normal hatching by decreasing intra-chorionic movement. A) Hatching rate measured from 0-4 dpf in DU-exposed and control larvae, displayed using a Kaplan-Meier survival analysis, with hatching on the y-axis and treatment across the x-axis; significance was determined by Logrank (Mantel - Cox) test and Fisher’s exact test (*, p < 0.05). B) Cathepsin L ELISA from DU-exposed and control larvae, analyzed by Kruskal-Wallis test and Dunn’s Test for Multiple Comparisons. C) Intra-chorionic movement was evaluated during the critical hatching window between 72 and 83 hpf. The trial started with six larvae per treatment; by 84 hpf all control larvae were hatched. Outliers were removed when their mean value is outside the column Mean ± 2xSD. Using repeated measures, two-way ANOVA and Dunnett’s post test showed significantly reduced intra-chorionic movement by Dose (p<0.0001; F (3, 20) = 16.68), increased chorionic movement by Time (p=0.0179; F (4.213, 59.75) = 3.178), but there was no interaction of Dose x Time (p=0.4743; F (12.64, 59.75) = 0.9863). A single pairwise comparison from the 73 hpf timepoint was significant, (0 v 0.1, p = 0.0227). The graph displays the mean movement and SEM over the12 hour time course of the critical hatching window.

### 3.2 Depleted uranium particles cause proximity-dependent ultrastructural defects to mitochondria

To determine if localized exposure to triuranium octoxide impacts adjacent tissues, we used a DU shrapnel model in zebrafish larvae in conjunction with electron microscopy (Fig. 2A-C). Ultrastructural ranking reduced in proximal samples (median = 2.750) compared to sham (median = 4.167) or contralateral (median = 3.917) samples, suggesting a proximity-dependent effect at implant sites, but not in sham or contralateral sites (Fig. 2D).

**FIGURE 2.**
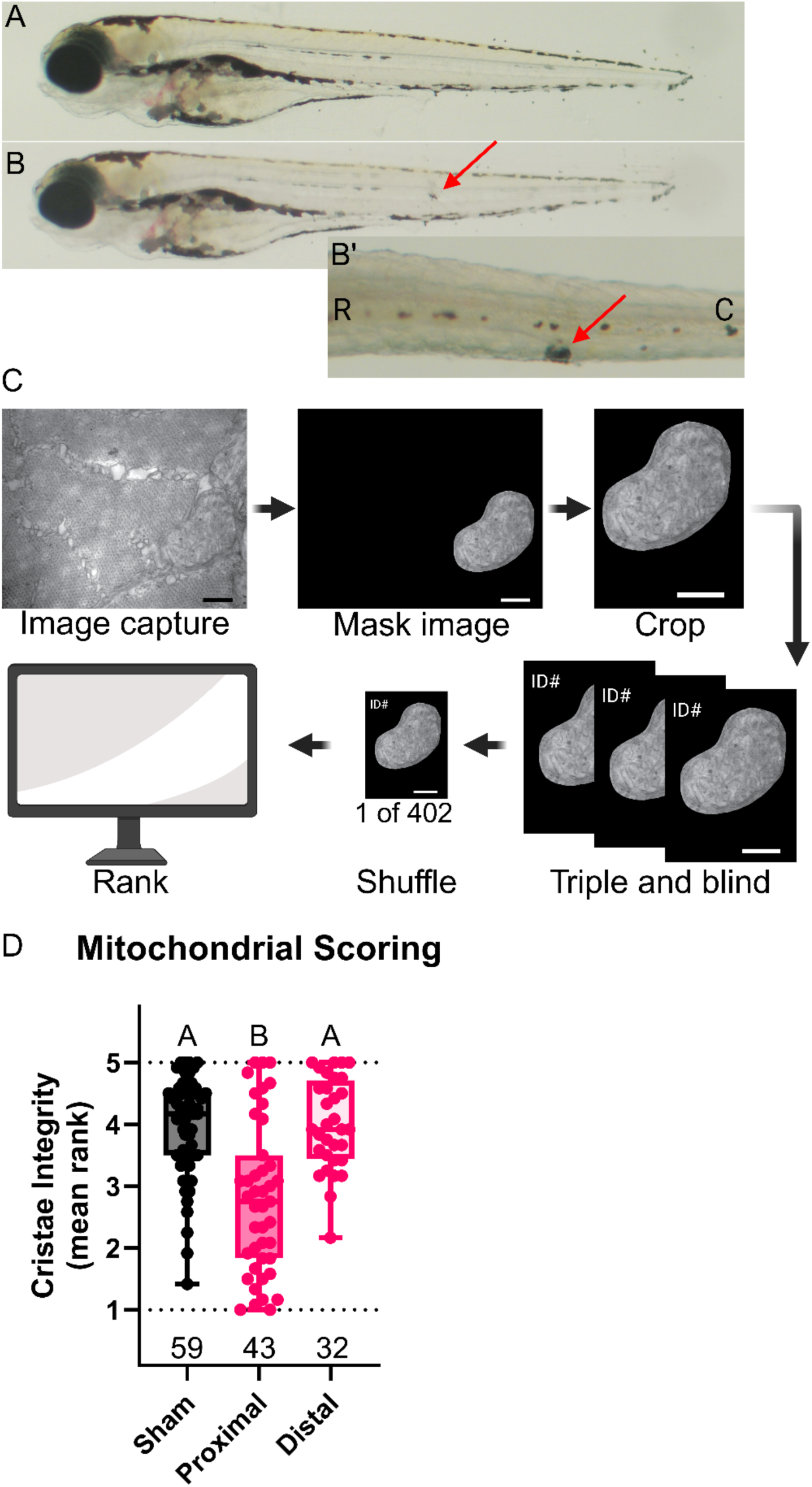
Depleted uranium particles cause proximity-dependent ultrastructural defects to mitochondria. A, B, B’) example light micrographs of U_3_O_8_ implanted larvae. A) lateral pre-, B) lateral post-implant view, red arrowhead denotes U_3_O_8_ implantation site, B’) post-implant from the dorsal view, R = rostral, C = caudal, red arrowhead denotes U_3_O_8_ implantation site on the left side of the animal. C) workflow for mitochondrial ranking. D) median ± SEM ranks of cristae integrity shown in a box and whiskers plot with n values (number of scored mitochondria) from three sham fish and three implanted fish (Kruskal-Wallis and Dunn’s, p < 0.05).

### 3.3 Overall reductive capacity is diminished by DU exposure and is unaffected by Nrf2 activation and targeted antioxidants

Cells with active mitochondria engaged in oxidative phosphorylation generate primary reducing molecules (*e.g.,* NADH, FADH_2_). The reagent alamarBlue^TM^ (ThermoFisher) measures cytosolic reductive potential by monitoring the change in fluorescence as the dye changes from the weakly fluorescent resazurin to the strongly fluorescent resorufin (Fig. 3A). Assay sensitivity was confirmed by a cell-free assay using increasing amounts of dithiothreitol (DTT) that produced a dose-dependent increase in alamarBlue signal (Suppl. Fig. 1). Larvae and PBMCs treated with increasing doses of DU showed a dose-dependent decrease in alamarBlue fluorescence, indicative of reduced mitochondrial activity (Fig. 3C-D), and quantitative real-time PCR revealed dose-dependent decreases in expression of antioxidant-response element (ARE) regulated genes *gss*, *gstp*, and *nqo1* (Fig. 3E-G). To test if reductive capacity can be restored by activation of ARE-regulated genes via its canonical activating transcription factor, Nrf2, embryos were pre-treated from 4 – 24 hpf with sulforaphane (SFN), a Nrf2 activator, then co-exposed with DU or As with continued addition of SFN. As expected, DU- and As-exposed larvae showed diminished reductive capacity; however, As-related decreases in reductive capacity were rescued by the presence of SFN, whereas DU-treated larvae showed no protection from SFN (Fig. 3H) despite canonical Nrf2 target activation as evidenced by an approximate 3-fold enrichment of *gstp* in all SFN-treated larvae, independent of uranium exposure (Fig. 3J). HEK 293 cells also showed that SFN rescued the metabolic defects imparted on As- but not DU-exposed cells (Fig. 3I). Rescue attempts utilizing the cytosolic antioxidant TEMPOL and the mitochondrially-targeted variant MitoTEMPO revealed that both drugs were unable to rescue DU’s metabolic deficits (Fig. 3K, L).

**FIGURE 3.**
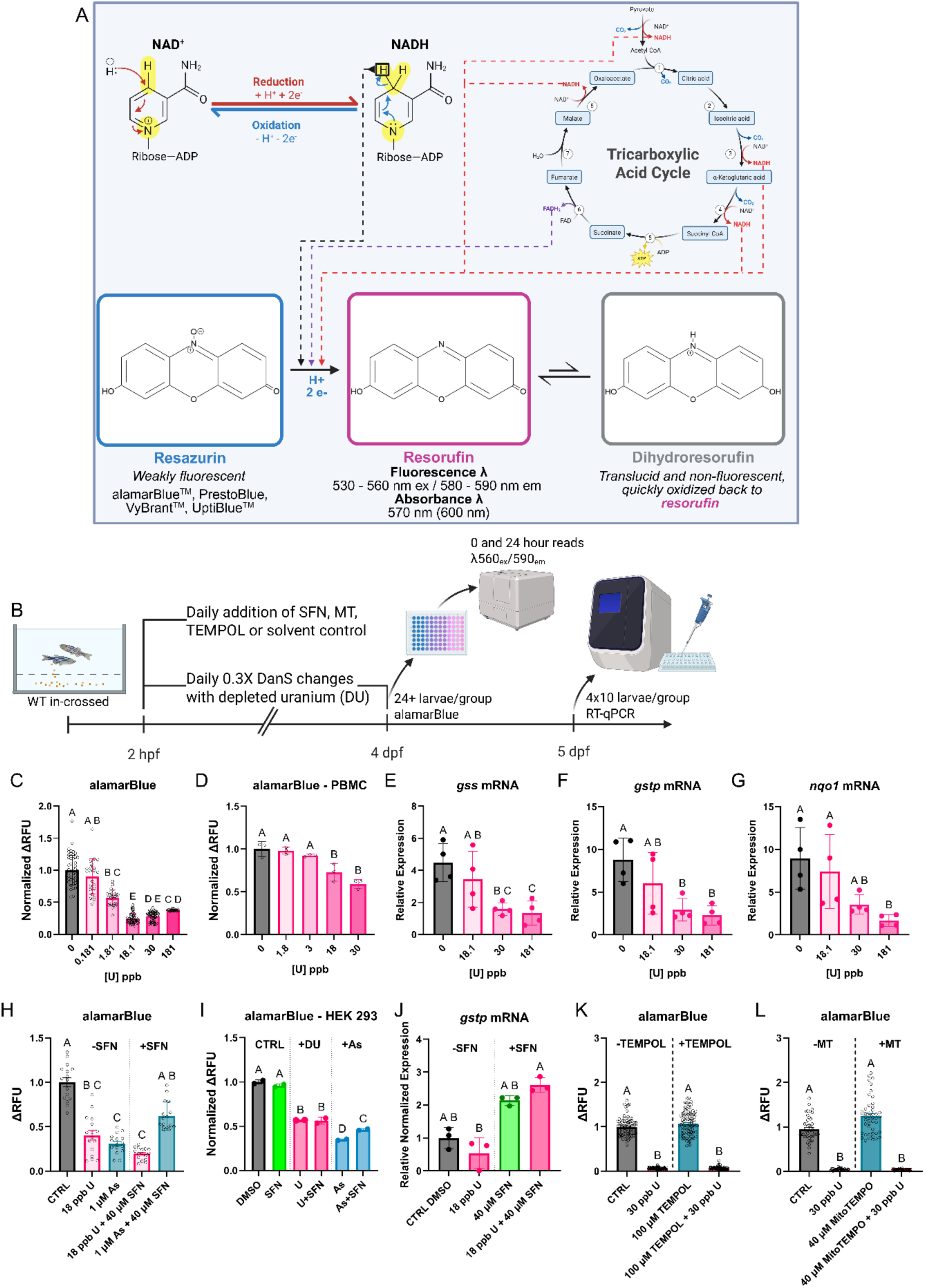
Overall reductive capacity is diminished by DU exposure and is unaffected by Nrf2 activation and targeted antioxidants. A) Overall chemical mechanisms underlying the alamarBlue assay. B) Experimental flowchart demonstrating waterborne DU experiments in addition to sulforaphane (SFN) + and antioxidant (TEMPOL/MitoTEMPO (MT)) challenges. C) Change in alamarBlue fluorescence over 24 hours in response to increasing doses of DU, total n = 227 individual zebrafish larvae and representative of three independent experiments across two separate genotypes (AB and Tü), analyzed using Kruskal – Wallis test and Dunn’s test for multiple comparisons, letters denote statistical significance (p < 0.05). D) Change in alamarBlue fluorescence over 20 hours in response to a dose curve of DU in PBMCs, n = three wells per group, analyzed using One-Way ANOVA followed by Tukey’s post-hoc multiple comparisons test, letters denote statistical significance (p < 0.05). E-G) Relative mRNA expression measured by RT-qPCR for ARE-regulated genes in zebrafish for E) *gss*, F) *gstp*, and G) *nqo1,* n = 4 biological replicates of ten pooled 5 dpf zebrafish larvae, analyzed using lognormal one-way ANOVAs with Tukey’s post-hoc multiple comparisons test, letters denote statistical significance (p < 0.05). H) Change in alamarBlue fluorescence over 24 hours in response to sub-MCL DU (18 ppb U) and MCL As (1 µM), in addition to 40 µM sulforaphane (SFN) rescue attempt, total n = 80 individual 5 dpf larvae, analyzed using Kruskal – Wallis test and Dunn’s test for multiple comparisons, letters denote statistical significance (p < 0.05); I) Change in alamarBlue fluorescence over 2 hours in HEK293T cells, analyzed using lognormal One-Way ANOVA with Tukey’s post-hoc multiple comparisons test, n = 2 seeded wells per group. J) Relative normalized expression of zebrafish *gstp* mRNA in response to sub-MCL DU with and without 40 µM SFN treatment; n = 3 biological replicates of ten 5 dpf zebrafish larvae, analyzed using an ordinary one-way ANOVA with Tukey’s post-hoc multiple comparisons test, letters denote statistical significance (p < 0.05). K) Change in alamarBlue fluorescence over 24 hours in response to MCL DU (30 ppb U) with and without 100 µM TEMPOL, total n = 334 individual 5 dpf larvae, analyzed by Kruskal – Wallis test and Dunn’s test for multiple comparisons, letters denote statistical significance (p < 0.05). L) Change in alamarBlue fluorescence over 24 hours in response to MCL DU (30 ppb U) with and without 40 µM MitoTEMPO, total n = 184 individual 5 dpf larvae, analyzed by Kruskal – Wallis test and Dunn’s test for multiple comparisons, letters denote statistical significance (p < 0.05).

### 3.4 Depleted uranium exposure affects global, nuclear, and mitochondrial DNA damage and decreases mitochondrial DNA copy number at MCL levels

To determine the contribution of genotoxicity to organismal phenotypes, we employed the comet assay and semi-long run qPCR for DNA damage quantification (Fig. 4A, 4D). Zebrafish larvae exposed to DU showed increased in Olive Tail Moments (OTMs; an indicator of DNA damage calculated by fluorescence intensity, dispersion, and tail length) under alkaline conditions (Fig. 4B) but not neutral comet (Fig. 4C). Damage imparted by UV irradiation was quantified by SLR qPCR for three DNA compartments (1978 bp mtCDS, 1939 bp mtD-loop, 1637 bp nucCDS), producing EC_50_ values of 25.92, 27.62, and 26.58 mJ/cm^2^ respectively (Suppl. Fig. 2A-C). Damage measured at the a*hr* gene locus was approximately 3-fold higher in exposed versus unexposed larvae by SLR qPCR in one of two gene targets, with the other showing no significant trends (Fig. 4H-I, Suppl. Fig. 2D). Damage to the mitochondrial genome was less clear, occasionally significant damage accumulation or reductions, but often showed no significant changes in damage (Fig. 4E-G; Suppl. Fig. 2D). Mitochondrial genome copy number (mtDNAcn) was diminished in exposed larvae compared to control, regardless of uranium dose (Fig. 4J) and across several experiments (Suppl. Fig. 2D). DU exposure also caused a stepped decrease in citrate synthase specific activity (Suppl. Fig. 3).

**FIGURE 4.**
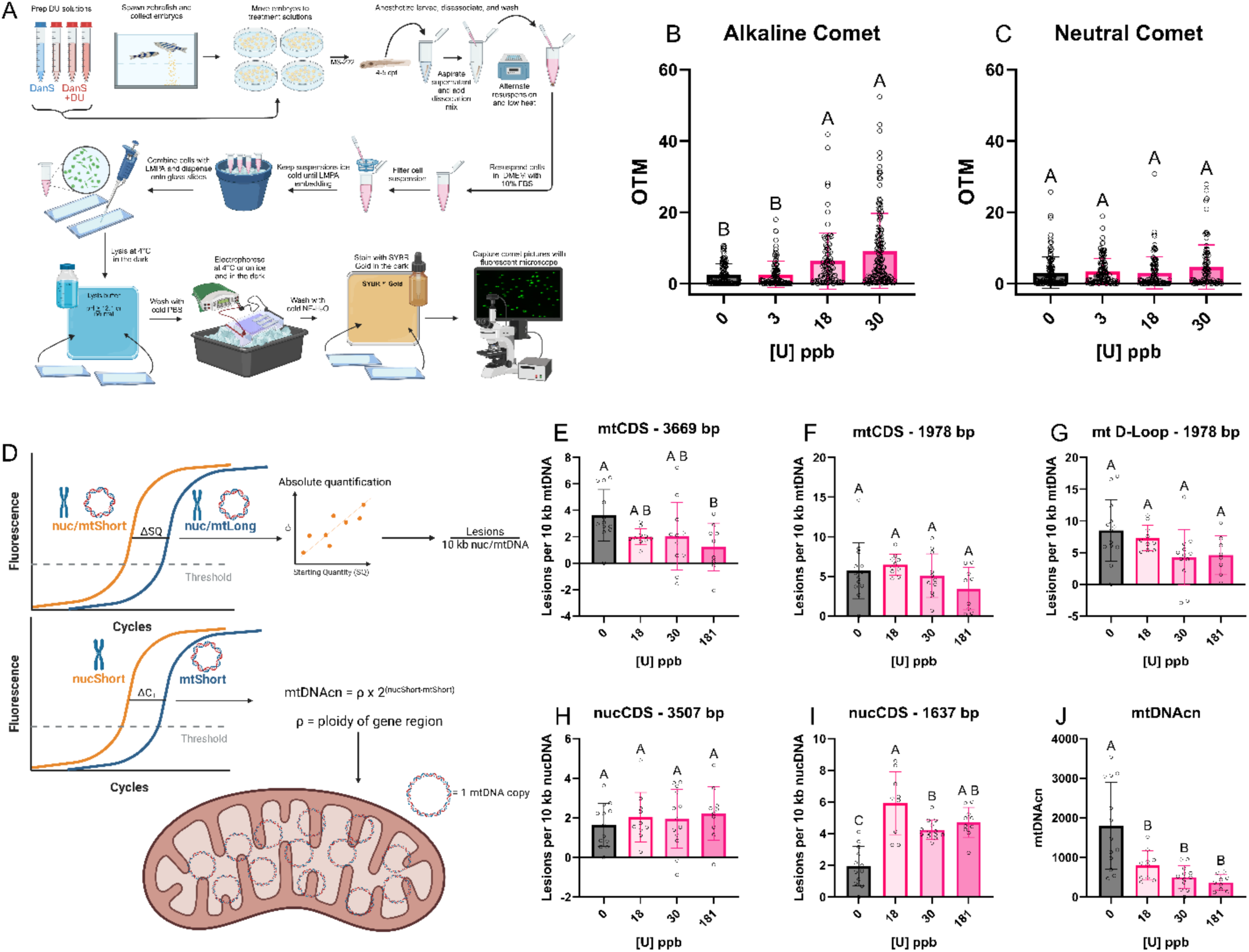
Depleted uranium exposure modulates global and nuclear, but not mitochondrial DNA damage, and decreases mtDNA copy number at sub-MCL levels. A) Experimental outline for single-cell gel electrophoresis (Comet) assay for detecting global DNA damage. B) Alkaline and C) neutral comet Olive tail moments (OTMs) blinded and scored, analyzed using Kruskal – Wallis test and Dunn’s test for multiple comparisons, n = 3-4 disassociated 5 dpf zebrafish embryos. D) SLR qPCR experimental diagram depicting compartment-specific lesion frequency calculations (top) and mitochondrial DNA copy number (mtDNAcn) determination. E-J) SLR qPCR lesion and mtDNAcn analyses. E) mitochondrial DNA coding sequence, 3669 bp amplicon length, F) mitochondrial DNA coding sequence, 1978 bp amplicon length, G) mitochondrial DNA D-loop, 1939 bp amplicon length, H) nuclear *ahr1* locus, 3507 bp amplicon length, I) nuclear *ahr1* locus, 1637 bp amplicon length, lesion frequency per 10 kilobase, analyzed by One-Way ANOVA and Holm-Šídák’s multiple comparisons test for multiple comparisons, letters denote statistical significance (p < 0.05). J) mtDNAcn, analyzed by One-Way ANOVA and Holm-Šídák’s multiple comparisons test, letters denote statistical significance (p < 0.05).

### 3.5 RNA sequencing and RT-qPCR reveal dose-dependent enrichment of *ssh1b* in response to DU

To discover differentially expressed gene targets of uranium, RNA sequencing was conducted on pooled 5 dpf zebrafish larvae (n = 4, where each sample is a pool of ten larvae; Fig. 5A). 181 ppb U did not show enrichment of Nrf2-target genes but did reveal 22.4-fold enrichment of *ssh1* mRNA by RNA-sequencing, the most significantly upregulated gene target in the dataset (Fig. 5B). Quantitative PCR for the four isoforms of the *ssh1* gene expressed by zebrafish reveals that *ssh1b* is dose-dependently enriched by up to 18-fold in response to DU exposure, while all other isoforms did not respond (Fig. 5C-F; Suppl. Table 1).

**FIGURE 5.**
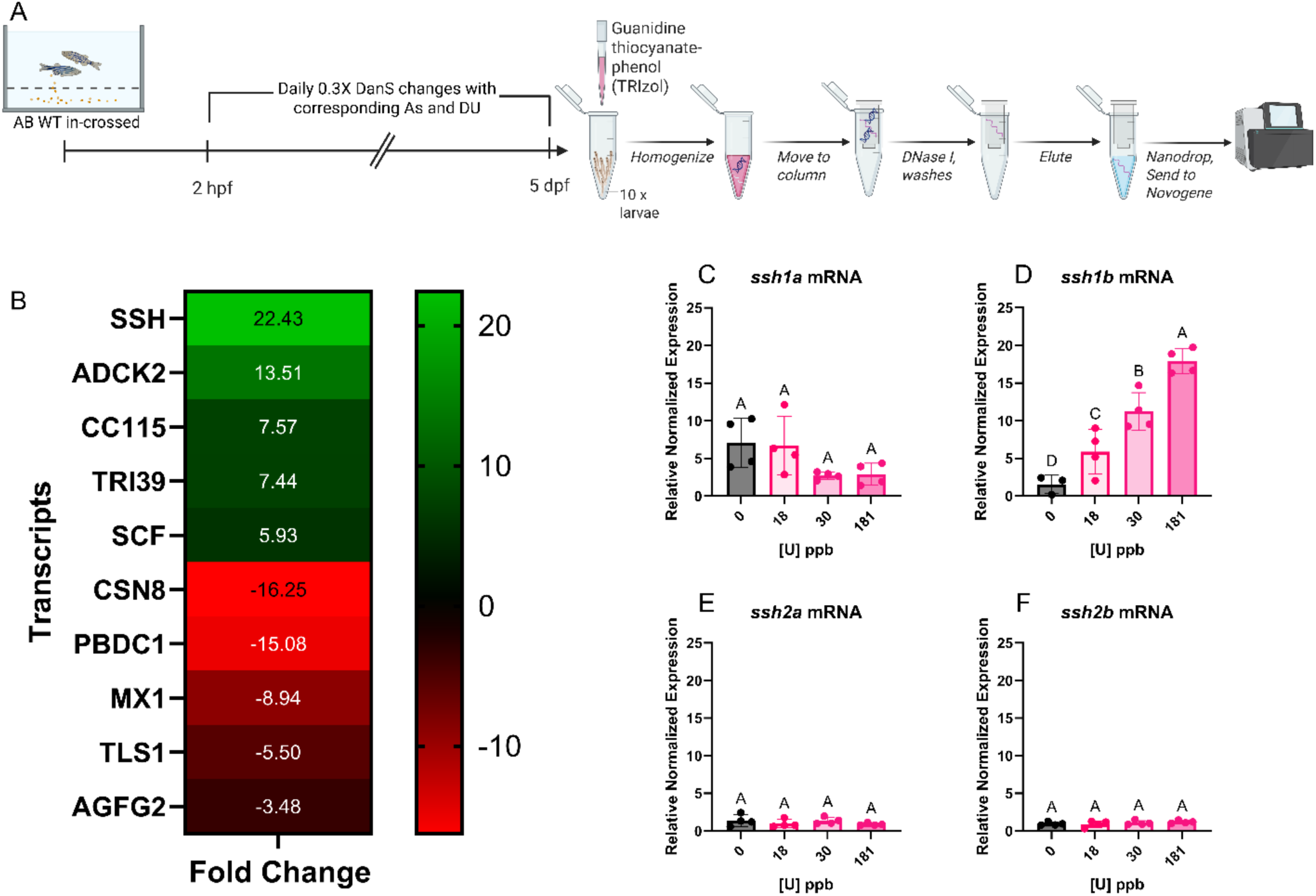
RNA sequencing and RT-qPCR reveal dose-dependent enrichment of ssh1b in response to DU. A) Experimental flowchart for RNA-sequencing experiment. B) Fold change of the top 5 up- and down-regulated differentially expressed genes (DEGs) for 181 ppb U vs. vehicle control. C-F) Relative normalized expression of all zebrafish *ssh* orthologs, n = 3 – 4 biological replicates of ten pooled 5 dpf larvae; from top left to top right: C) *ssh1a* and D) *ssh1b* mRNA, from bottom left to bottom right: E) *ssh2a* and F) *ssh2b* mRNA. *ssh1a*, *ssh2a*, and *ssh2b* were all analyzed using lognormal ordinary one-way ANOVAs followed by Tukey’s multiple comparisons test while *ssh1b* expression was analyzed using ordinary one-way ANOVA followed by Holm-Šídák’s multiple comparisons test, letters denote statistical significance (p < 0.05).

### 3.6 Sennoside A can rescue DU-induced metabolic output deficiencies and hatching defects

We attempted a rescue experiment through inhibition of the Ssh family of proteins using previously reported *in vitro* dosing of 10 µM Sennoside A (Fig. 6A). Metabolic analyses via alamarBlue revealed that 10 µM Sennoside A could ameliorate DU-induced metabolic defects (Fig. 6B). Comparisons of hatching rates revealed that 10 µM Sennoside A was also able to restore normal hatching rates in the presence of sub-MCL DU (Fig. 6C, D). Figure 6E illustrates the molecular hypotheses underlying the experiment displayed in Fig. 6F. Metabolic analyses via alamarBlue also revealed that SFN administration in the presence of 10 µM Sennoside A in the context of DU exposure increases metabolism compared to the DU + SA treatment group, while additions of Brusatol, MitoTEMPO, and TEMPOL in the presence of DU + SA did not modulate metabolism compared to DU + SA alone (Fig. 6F).

**FIGURE 6.**
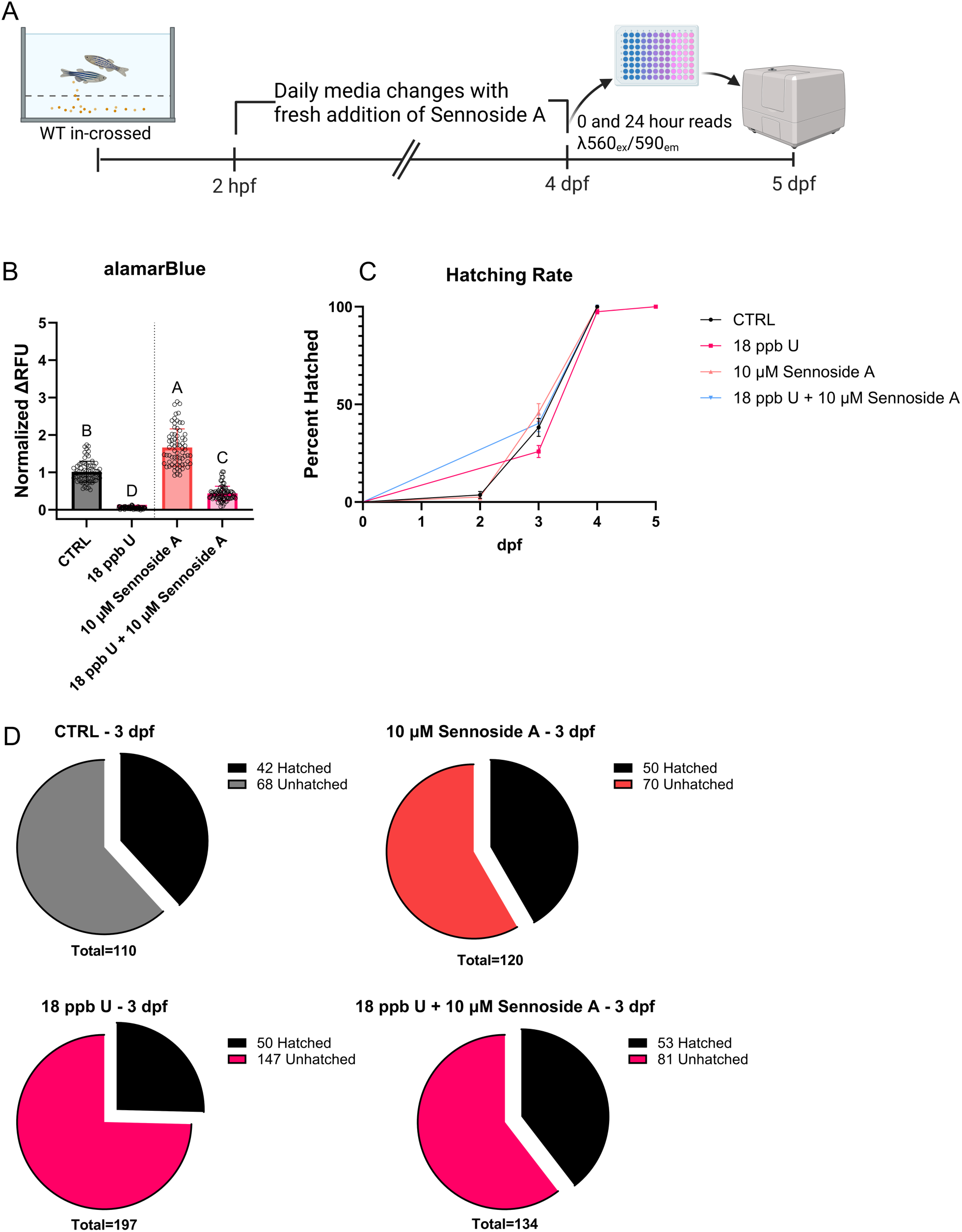
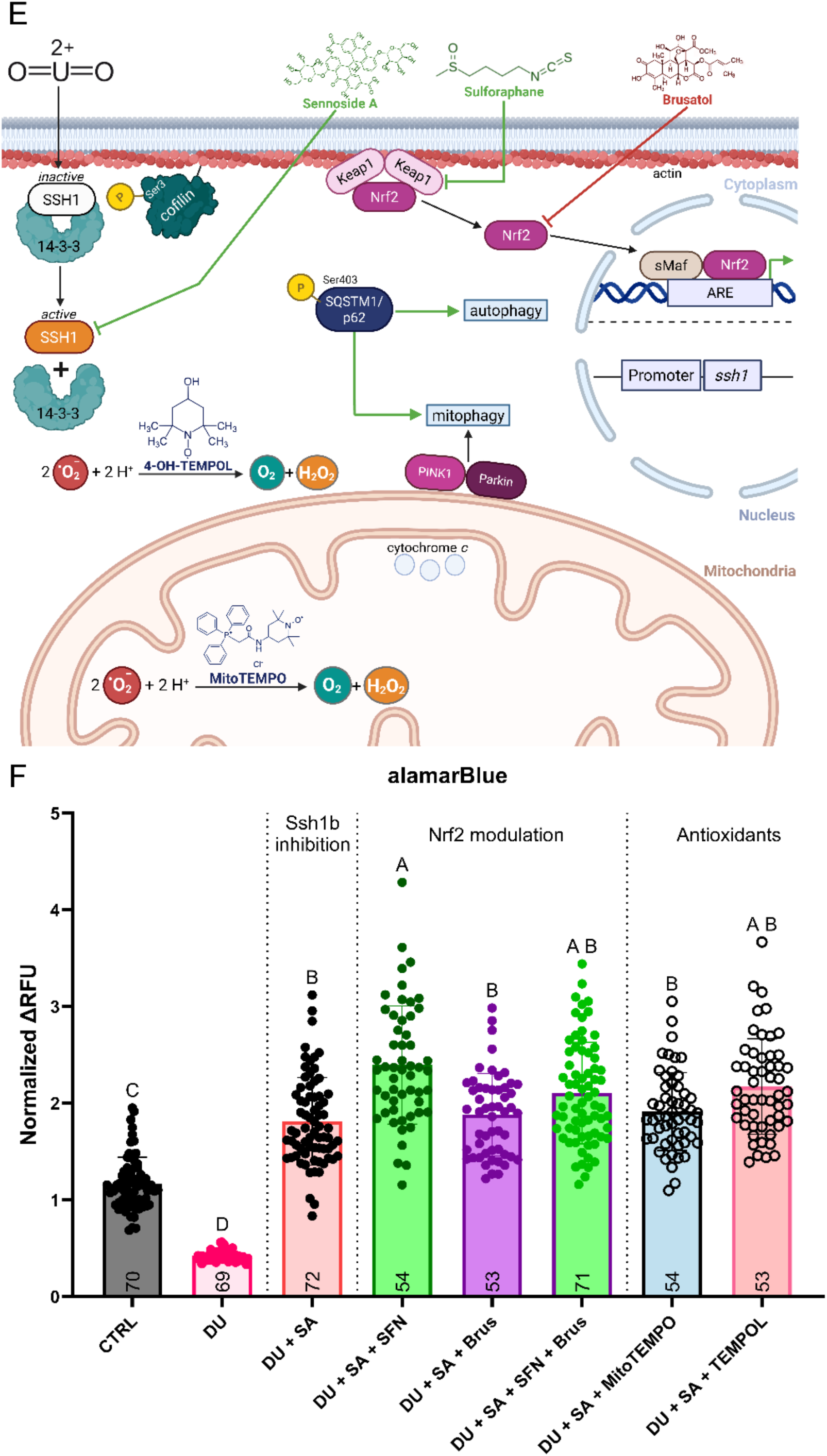
Sennoside A can rescue DU-induced metabolic output deficiencies and hatching defects. A) Experimental flowchart for daily Sennoside A administration and subsequent alamarBlue assay. B) Change in alamarBlue fluorescence of individual zebrafish larvae over 24 hours in response to sub-MCL (18 ppb) DU, with and without daily addition of 10 µM Sennoside A, analyzed using Kruskal – Wallis test and Dunn’s test for multiple comparisons and data are representative of two independent experiments in AB genotype zebrafish, total n = 300 individual 5 dpf larvae. C) Hatching rate measured from 0 – 5 dpf in treated larvae, displayed using a Kaplan-Meier survival analysis; significance was determined by log-rank (Mantel - Cox) test for the overall Kaplan-Meier analysis (p = 0.0002). D) Parts-of-whole depictions of 3 dpf hatching rates shown in C), analyzed using Fisher’s exact test (p = 0.0061). E) BioRender diagram depicting the hypothesized effects and interactions of Sennoside A (SA), Sulforaphane (SFN), Brusatol (Brus), 4-OH-TEMPOL (TEMPOL), and MitoTEMPO (MT) on intracellular dynamics in the presence of the uranyl ion (UO_2_^2+^). F) Change in alamarBlue fluorescence of individual zebrafish larvae over 24 hours in response to 18 ppb DU and 10 µM Sennoside A (SA) alone and in combination with Nrf2 modulating drugs (40 µM sulforaphane (SFN) and 500 nM Brusatol (Brus)) both alone and in combination, and compartment-specific antioxidants alone (40 µM MitoTEMPO, 100 µM TEMPOL). Data were analyzed by Kruskal - Wallis test and Dunn’s test for multiple comparisons, representative of one independent experiment in AB genotype zebrafish, total n = 496 individual 5 dpf larvae, n value per group is annotated at the bottom of each bar (n = 53 – 72 per group).

## 4. DISCUSSION

In this report, we set out to determine the mechanisms underlying the chemical toxicity of depleted uranium. Leading hypotheses in the field all posit that DU causes: 1) ROS generation causing oxidative stress and potential mitochondrial damage as a consequence, 2) DNA damage accumulation producing genotoxicity, 3) protein binding producing abnormal function (transferrin, ferritin, fetuin-A), 4) inflammation activation, and 5) apoptosis and autophagy activation (Ma et al., 2020).

### 4.1 Gulf War Illness

Gulf War Illness (GWI) is an idiopathic chronic illness that affects Gulf War veterans (Elhaj and Reynolds, 2023). One of the most widely claimed symptoms of Gulf War Illness, as defined in the Kansas Study, is fatigue (Steele, 2000). The American Gulf War I saw extensive implementation of DU-containing munitions (Fetter and Von Hippel, 2000; Jargin, 2014). There has been substantial research on the topic; however, GWI’s cause remains mysterious. Amplifying the mystery are conflicting reports; some suggest DU-exposure is a potential cause (Bjørklund et al., 2020; Ma et al., 2020) while others refute that interpretation (Parrish and Haley, 2021). Studies using PBMCs from Gulf veterans showed depressed bioenergetics, but the study did not look at uranium exposure specifically (Meyer et al., 2023). Parrish and Haley’s study used ultrasensitive mass spectroscopy and bioassays to examine cell health in Gulf War veterans and showed that uranium was not detectable despite patients being symptomatic. They conclude that this is proof of uranium not contributing to the GWI state; however, they do not consider if uranium exposure sensitized subjects leading to phenotypes that persist long after exposure has ceased (Parrish and Haley, 2021). Additionally, it seems possible the GWI could manifest because of different exposure paradigms and/or chemicals. That is to say that DU could contribute, but it is not by itself causative.

### 4.2 Depleted uranium delays normal hatching by decreasing intra-chorionic movement

The only reliable whole organism phenotype we observed in depleted uranium exposed larvae was delayed hatching by approximately 24 hours (Fig. 1A), attributed to reduced embryonic movement in the chorion (Fig. 1C), after determining there was no change in the release of the primary hatching enzyme, Cathepsin L (Fig. 1B). Reduced movement in the chorion prior to hatching may be explained by mitochondrial injury, genotoxicity, ROS accumulation, or any other host of energetic-dampening perturbant.

### 4.3 Depleted uranium particles cause proximity-dependent ultrastructural defects to mitochondria

To determine if localized exposure to triuranium octoxide impacts adjacent tissues, we used our zebrafish DU shrapnel model and electron microscopy (Fig. 2A-C). Micrographs of mitochondria from the sham site, implant site, or contralateral to implant site were blinded, ranked, unblinded, and resulting data were analyzed. Ultrastructural ranking was significantly reduced, with the median score of 2.750 for proximal sites, and 4.167 and 3.917 for sham and contralateral sites, respectively (Fig. 2D). We take these data to mean that DU imparts a proximity-dependent effect on mitochondrial cristae ultrastructure, that remains limited to only the implanted side of the organism. This also informs us that retained DU fragments in humans (like those from McDiarmid and company’s Gulf War cohort studies) may negatively impact mitochondrial health in tissues proximal to the fragment, but tissues located further away from a fragment are likely to remain relatively healthy since the effect was only observed at the implant site, indicating minimal systemic mobilization of the particles.

### 4.4 Overall reductive capacity is diminished by DU exposure that is unaffected by Nrf2 activation and targeted antioxidants

To determine the contribution of ROS and REDOX-related processes in uranium toxicity, we exposed zebrafish and cells to varying doses of DU or DU with an antioxidant challenge (Fig. 3B). DU decreased the reductive capacity of both PBMCs and zebrafish larvae in a dose-dependent manner at doses below the EPA MCL (Fig. 3C-D). One main hypothesis in the field is that uranium modulates dynamics of oxidative stress and depletes the antioxidant response through increases in interactions with thiobarbituric acid-reactive substances (TBARS), GSH, SOD, and GR expression (Bellés et al., 2005) in addition to an increase in the activity of SOD, catalase, and GPx (Lestaevel et al., 2009; Ma et al., 2020). We similarly demonstrate that DU decreases relative mRNA expression of three ARE-regulated genes in zebrafish, *gss*, *gstp*, and *nqo1* (Fig. 3E-G). Arsenic has been shown to increase intracellular ROS levels, activating the Nrf2-Keap1 pathway to detoxify these ROS from the cell (Lau et al., 2013). Therefore, we hypothesized that since Nrf2 activators have been shown to ameliorate or prevent As-induced increases in ROS (Hu et al., 2020), and if DU proceeds through a ROS-dependent mechanism, then a Nrf2 activator would have a similar result. We first tested whether sulforaphane (SFN), a potent Nrf2 activator, could rescue basal metabolic activity from metal-produced ROS. We showed that SFN was able to increase the metabolic activity of As-treated larvae, but not DU-treated larvae (Fig. 3H), despite SFN-induced transcription of *gss* in DU-treated larvae (Fig. 3J). This relationship was consistent in HEK 293 cells exposed to DU and As with or without SFN, where SFN rescued As- but not DU-induced defects (Fig. 3I). We also demonstrated that two SOD-mimetic compounds, TEMPOL and MitoTEMPO were both unable to ameliorate DU-induced metabolic deficits (Fig. 3K, L). Taken together, these results demonstrate that DU toxicity must proceed by way of a mechanism independent of either Nrf2 activation or ROS-production. That is not to say that Nrf2 is not involved in the toxicity of uranium or its’ effects as we demonstrate dose-dependent decreases in ARE-regulated genes; however, our results also show that in the context of larval uranium exposure, Nrf2 induction using SFN does not modulate metabolic state, despite increased expression of ARE-regulated genes, suggesting that ROS-related impacts on mitochondrial function is not the only mechanism of uranium toxicity.

### 4.5 Depleted uranium exposure modulates global, nuclear, and mitochondrial DNA damage and decreases mitochondrial DNA copy number at MCL levels

One leading hypothesis for depleted uranium chemical toxicity is that uranyl ions adduct to the phosphate backbone of DNA, potentiating strand breakage (Yazzie et al., 2003). We hypothesized that DNA damage and resulting genotoxicity derived from uranyl adduction to the phosphate backbone of DNA could drive uranium toxicity. We performed single-cell electrophoresis (*i.e.,* comet) for global DNA damage quantification (Fig. 4A) and semi-long run (SLR; Fig. 4D) quantitative PCR for compartment-specific DNA damage quantification on zebrafish larvae exposed to varying concentrations of DU (Zhu and Coffman, 2017). Comet assay for global DNA damage in disassociated zebrafish larvae produced increased alkaline comet OTM but no significant increase in neutral comet OTM suggesting that DU causes lesions and/or labile sites that are converted to DSB under alkaline comet conditions, but not neutral conditions (Fig. 4B-C) (Pribakovic et al., 2025). SLR qPCR on exposed 5 dpf zebrafish larvae revealed that nuclear DNA damage accumulated in one gene target, but not the other, raising questions as to the stochasticity of uranium-DNA adduction in the nucleus (Fig. 4H, I, and Suppl. Fig. 2D). Changes in mitochondrial DNA damage were detected, but the changes were inconsistent across experiments and target regions with increases and decreases observed (Fig. 4E-G, and Suppl. Fig. 2D); we therefore conclude that mitochondrial DNA does not appear to be a target of uranium-induced DNA damage. Mitochondrial DNA copy number (mtDNAcn) consistently decreased in the presence of uranium at any concentration (Fig. 4J and Suppl. Fig. 2D). Aligning with this observation, we show that DU decreases citrate synthase specific activity in a stepped manner, interpreted as a decrease in mitochondrial mass (Chhimpa et al., 2023; Gillen et al., 2016; Liu et al., 2018), suggesting a more complex mitochondrial mechanism at play. These data support the hypothesis that DU adduction to DNA potentiates DSBs but is not sufficient to cause DSBs independently (Stearns et al., 2005), while also dose-dependently decreasing mtDNAcn. Decreased mtDNAcn can be interpreted multiple ways – either there are less mitochondria featuring a similar number of copies to control or there are less copies per mitochondrion, with mitochondrial number unchanged. Taken with the finding of a stepped reduction in CS specific activity, we present two pieces of evidence for decreases in mitochondrial mass. This work also provides evidence for compartment-specific DNA damage, where nuclear but not mitochondrial damage was noted in response to DU. Altogether, these data confirm that uranium is a genotoxicant *in vivo*, where the nuclear genome is a target, but hints at a more complex mitochondrial mechanism.

### 4.6 RNA sequencing and RT-qPCR reveal dose-dependent enrichment of *ssh1b* in response to DU

We designed an RNA-sequencing experiment to identify novel gene targets of uranium toxicity (Kalaniopio et al., 2026; Fig. 5A). The most significantly up-regulated gene in response to 181 ppb U was mapped to SSH, the slingshot homolog family of genes (Fig. 5B). SSH1 is implicated in mitochondrial health and mitophagy/autophagy in human cells through novel N-and C-terminal domains that interact with cofilin and p62 respectively (Cazzaro et al., 2023b) while also sequestering Nrf2, preventing activation of detoxifying enzymes (Cazzaro et al., 2023a). After identification of SSH ortholog as a regulator of mitophagy in the literature, we performed targeted real-time PCR to identify which of four possible zebrafish-specific paralogs were expressed in response to uranium exposure – revealing *ssh1b* as the only dose-dependent responsive gene in the family of expressed transcripts (Fig. 5C-F) aligning with conclusions from the literature. Taken together, we show that *Ssh1b* is highly responsive to uranium exposure in zebrafish, indicating that it is a primary target of toxicity and the modulator of mitochondrial phenotypes, but a full gain- and loss-of-function exploration is required.

### 4.7 Sennoside A partially rescues DU-induced metabolic output deficiencies and hatching defects

Sennoside A (SA), a dianthrone glycoside derived from the senna family and used clinically as a laxative, has been shown to be a potent inhibitor of the functions of the slingshot homolog family of proteins in human cell lines, proceeding through an unknown mechanism (Lee et al., 2017). We hypothesized that inhibition of the SSH family at the protein level using SA would reduce uranium’s effects on the mitochondria through reduced Ssh1b activity and thereby a lack of inhibition of mitophagy, leading to increased overall metabolic activity, rescue of hatching rate disruptions, and an overall amelioration of uranium toxicity by available metrics. Metabolic analysis by alamarBlue revealed that 10 µM SA was able to recover uranium-derived metabolic defects (Fig. 6B). This partial recovery of metabolic activity proved sufficient to also recover the hatching rates of exposed larvae to near that of control (Fig. 6C-D). In the PBMCs of veterans with Gulf War Illness, decreased energetics through some form of mitochondrial insult was noted (Meyer et al., 2023). Furthermore, we hypothesized that inhibition of Ssh1b (or SSH1) with SA would allow Nrf2 modulators like SFN and Brusatol (Brus) or antioxidants like MitoTEMPO and TEMPOL to exert their canonical function, ultimately modulating alamarBlue-detectable metabolic changes (Fig. 6E). Indeed, addition of 10 µM SA in the presence of 18 ppb DU created conditions in which Nrf2 was most likely not sequestered by SSH and therefore available for chemical manipulation, as demonstrated by increases in fluorescence of the DU + SA + SFN group compared to the DU or DU + SA groups alone (Fig. 6F). Surprisingly, effective concentrations of Brusatol (500 nM, a potent Nrf2 inhibitor), MitoTEMPO (40 µM, mitochondrially targeted TEMPOL) and TEMPOL (100 µM, free-radical scavenging nitroxide) did not significantly change alamarBlue fluorescence in the presence of DU + SA (Fig. 6F). These data demonstrate that aberrant SSH signaling can be attenuated using Sennoside A in the context of uranium-driven SSH over-expression and over-activation *in vivo*. Furthermore, Nrf2 appears to be repressed by SSH activation in the context of uranium toxicity, which can also be ameliorated using SA, explaining why a successful rescue of metabolic function using SFN only occurs in the presence of SA (Fig. 6F), but not by itself (Fig. 3H-I). These findings support a mechanism of toxicity in which uranium exposure induces aberrant SSH signaling linked to mitochondrial damage, with minimal contributions from cellular ROS species or cellular redox responses. We propose that this damage is a consequence of impaired mitochondrial health and dysregulated mitophagy due to aberrant SSH signaling and its role as a modulator of autophagy and mitophagy that can explain phenotypes observed under environmentally relevant challenges of depleted uranium. *To our knowledge, this is the first evidence supporting Ssh1 as a biomarker for uranium chemical toxicity and a Molecular Initiating Event*.

## 5. CONCLUSION

Our results point to mitochondria as a cellular target of DU toxicity that leads to impaired mitochondrial structure and function, producing a REDOX imbalance. ROS has long been considered a driving factor of DU toxicity; however, inhibition of ROS accumulation using TEMPOL and MitoTEMPO at effective concentrations provided no protection, and activation of antioxidant response genes downstream of Nrf2 with SFN also had no effect. DNA damage is another leading mechanism for DU toxicity, and in agreement with previous studies, we show increased levels of nuclear but not mitochondrial DNA damage. Globally, we did not observe an increase in DSBs; however, increases in alkaline-labile sites were observed.

Gene expression studies following DU exposure pointed at dysregulation of the slingshot homolog (SSH) family of genes. In zebrafish we showed that DU exposure caused over-expression of *ssh1b,* and blocking Ssh1b activity using the only currently available SSH inhibitor, Sennoside A (SA), partially ameliorated DU’s toxic effects on metabolism and hatching. Furthermore, SA treatment enabled SFN to increase mitochondrial function above that of SA alone. These findings have large implications for Gulf War veterans exposed to DU, as well as for underserved populations with a chronic uranium burden, where future investigations should seek to identify human ortholog SSH1 as a biomarker and drug target for uranium exposure and subsequent toxicity.

## LIST OF SYMBOLS, ABBREVIATIONS AND FORMULAS

AB: AB-wildtype strain zebrafish
AP: armor-piercing
ARE: antioxidant response element
ATP: adenosine triphosphate
CRUMP: Church Rock Uranium Monitoring Project
DanS: Danieau’s solution
DTT: dithiothreitol
DMSO: dimethyl sulfoxide
dpf: days post-fertilization
DU: depleted uranium
EPA: Environmental Protection Agency
ETC: electron transport chain
EU: enriched uranium
FET: fish embryo toxicity test
Gss: glutathione synthetase
Gstp1: glutathione-S-transferase Pi
hr: hour
hpf: hours post fertilization
IAEA: International Atomic Energy Agency
IACUC: Institutional Animal Care and Use Committee
KEAP1: kelch-like ECH-associated protein 1
LOEL: Lowest observed effect level
MCL: Maximum Contamination Limit
Mn-SOD: manganese superoxide dismutase; SOD2
Mn-SOD: SOD2 Superoxide dismutase 2
mpf: months post-fertilization
mtDNA: mitochondrial
DNA NaAsO_2_: sodium meta arsenite
NAD+/NADH: nicotinamide adenosine dinucleotide
NOEL: No observed effect level
Nqo1: NAD(P)H dehydrogenase quinone 1 isoform
1NU: natural uranium
OPA1: optic atrophy-1
OXPHOS: oxidative phosphorylation
PGC-1α: peroxisome proliferator-activated receptor gamma coactivator 1-alpha ppb, parts per billion
ROS: Reactive Oxygen Species
RSF: resorufin
RZR: resazurin SFN sulforaphane
SOD2: manganese superoxide dismutase;
Mn-SOD Ssh1: slingshot protein phosphatase 1
TEM: transmission electron microscopy
TL: Tüpfel longfin strain zebrafish
TU: Tübingen strain zebrafish
U: uranium
UN: UO_2_(NO_3_)_2_, uranyl nitrate
v/v: volume to volume
w/v: weight to volume
WT: wildtype

## ACKNOWLEDGMENTS

This work was supported by a grant from the National Institutes of Environmental Health Sciences (R15ES032923-01/S1; MCS, PHK), a pilot grant from the Southwest Health Engangement Research Collaborative (U54MD012388; MCS) and student support from the National Science Foundation Louis Stokes Alliance for Minority Participation (HRD1712523; PHK), the Native American Partnership for Cancer Prevention (U54CA143925; PHK), Research Initiative for Scientific Engagement (1R25GM127199-01; RSA, ORL), Hooper Undergraduate Research Award (HLB, PHK), and the State of Arizona Technology and Research Initiative Fund (TRIF), administered by the Arizona Board of Regents. We are extremely grateful to our past and current undergraduate research assistants, Dr. Madisyn (Kaus) Peru, Hayden L. Bekkedahl, and Brooke Hinchcliff for contributing their time and effort toward optimizing experimental approaches. Lastly, we would like to thank Dr. Tracy Punshon and the entire Dartmouth BNEIR group for elemental analysis of exposed zebrafish, Dr. Jani Ingram for access to ICP-MS, and the NAU Office for Vice President for Research for access to microscopy and general research support.

## CRediT AUTHOR CONTRIBUTIONS

PHK: Conceptualization, data curation, formal analysis, funding acquisition, investigation, methodology, validation, visualization, writing – original draft, review and editing. MCS: Conceptualization, formal analysis, funding acquisition, methodology, visualization, writing – original draft, review and editing, resources, supervision, project administration. LBG, SMM ORL, EG, RSA: Conceptualization, data curation, formal analysis, investigation, methodology, validation, visualization, review. CAC, CMA, JW: data curation, formal analysis, investigation, methodology, validation, visualization, review. TT, CRP: writing – original draft, review, editing, supervision.

---

**SUPPLEMENTAL FIGURE 1**. alamarBlue four-point standard curve using dithiothreitol (DTT) in a cell-free system.

**SUPPLEMENTAL FIGURE 2.** SLR qPCR for DNA damage detection, compartmentalization, and trends analysis. A-C) SLR-qPCR standard curve of UVB irradiation on naked zebrafish DNA using A) 1978 bp mtCDS, B) 1939 bp mt D-loop, and C) 1637 bp nucCDS, R^2^ and EC_50_ values determined using GraphPad Prism. D) Trends sheets for all SLR qPCR experiments containing descriptive statistics and sample sizes with strain of zebrafish, p-value cells are color coded accordingly: red = ns, green = p < 0.05, yellow = 0.05<p<0.1.

**SUPPLEMENTAL FIGURE 3**. Citrate synthase specific activity measured across two independent experiments and two wildtype strains (AB and Tu), each dot represents protein extracted from n = 15-30 3 dpf zebrafish larvae. Analyzed by Kruskal-Wallis test and Dunn’s Test for Multiple Comparisons (p < 0.05).

**SUPPLEMENTAL FIGURE 4.** ICP-MS quantification of ^238^U concentrations in exposure solutions. Left column is reported dose of uranyl nitrate hexahydrate ((UO_2_(NO_3_)_2_) · 6 H_2_O) in micrograms per liter (ug/L) in zebrafish exposure solutions with relevant solvent controls (0.3x Danieau’s Solution, MilliQ-Water, and technical blank); middle column, theoretical ^238^U concentration in parts per billion (ppb) calculated from left column; right column, ^238^U concentration measured by ICP-MS in ppb.

**SUPPLEMENTAL TABLE 1**. Zebrafish RT-qPCR primers with direction, sequence, and source/citation.

**SUPPLEMENTAL TABLE 2**. SLR-qPCR primers for zebrafish mitochondrial and nuclear DNA targets, with validated parameters annotated for each primer set.

