## Supplemental materials for "Environmentally relevant depleted uranium exposure damages mitochondria, decreases cytosolic reductive capacity, and increases global DNA damage accumulation through a ROS-independent mechanism involving *slingshot protein phosphatase 1b* enrichment"

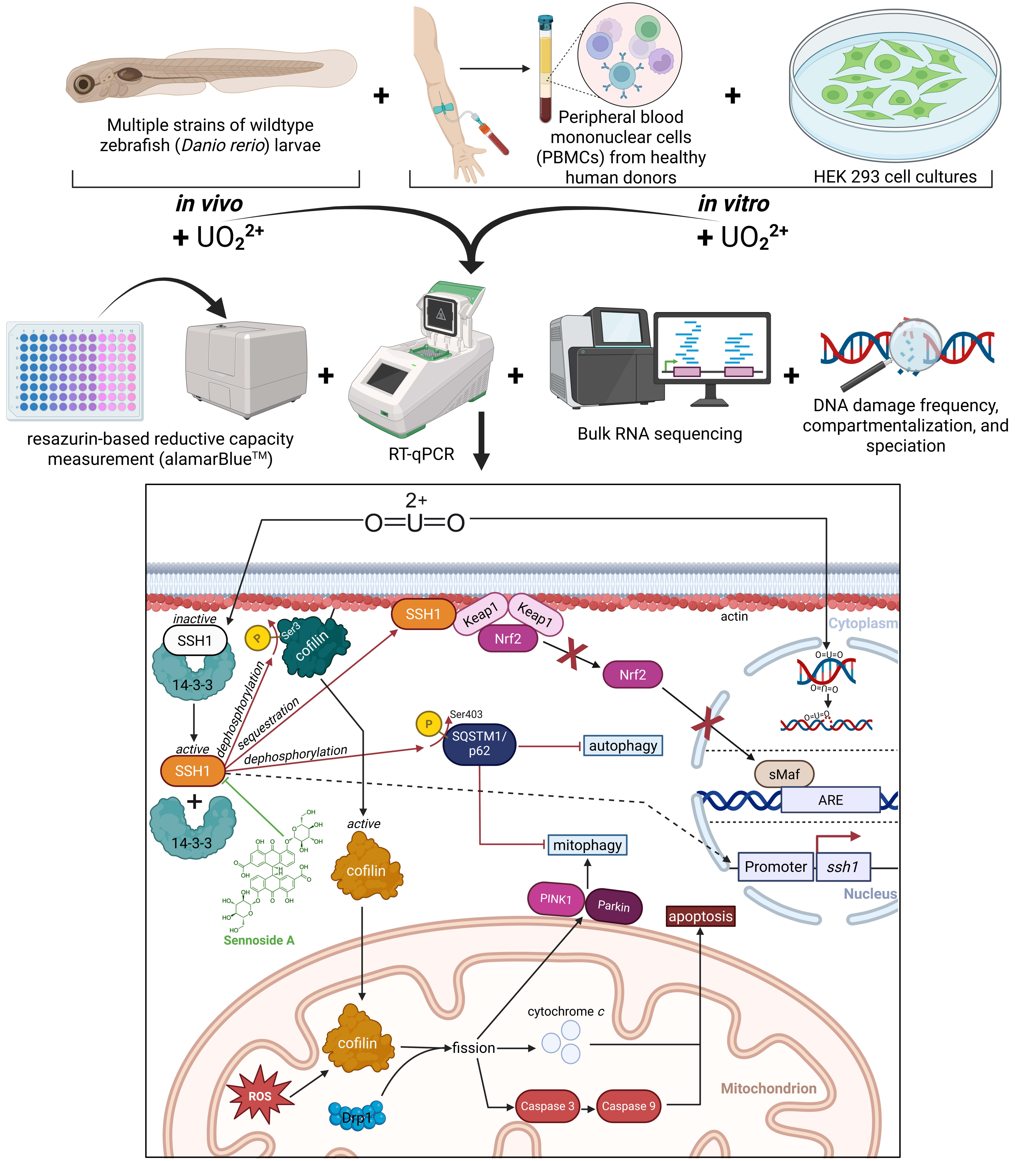

Multiple strains of wildtype zebrafish (*Danio rerio*) larvae
Peripheral blood mononuclear cells (PBMCs) from healthy human donors
HEK 293 cell cultures
*in vivo*
*in vitro*
+ $UO_2^{2+}$
+ $UO_2^{2+}$
resazurin-based reductive capacity measurement (alamarBlue™)
RT-qPCR
Bulk RNA sequencing
DNA damage frequency, compartmentalization, and speciation
2+
O=U=O
inactive
SSH1
14-3-3
P
Ser3
cofilin
SSH1
Keap1
Keap1
Nrf2
actin
Cytoplasm
Nrf2
dephosphorylation
sequestration
dephosphorylation
Ser403
SQSTM1/ p62
autophagy
active
SSH1
sMaf
ARE
14-3-3
active
cofilin
mitophagy
Promoter
*ssh1*
Nucleus
Sennoside A
PINK1
Parkin
apoptosis
cytochrome *c*
cofilin
fission
ROS
Drp1
Caspase 3
Caspase 9
Mitochondrion

**HIGHLIGHTS**

- Depleted uranium (DU, uranyl nitrate $UO_2(NO_3)_2$) decreases overall cellular-reducing capacity and increases DNA-damage accumulation in larval zebrafish and human cells at levels below the US EPA's MCL of 30 ppb U while implanted depleted uranium particles ($U_3O_8$) cause proximity dependent disruption of mitochondrial cristae structure.
- DU causes increased transcription of *ssh1b* and subsequent Ssh1b activation in zebrafish, driving mitochondrial and behavioral phenotypes.
- Inhibition of Ssh1b using Sennoside A recovers DU-induced metabolic and hatching rate defects in zebrafish.
- Sennoside A-driven inhibition of SSHs permits Nrf2 function using the Nrf2 activator, sulforaphane.
- Cytosolic and mitochondrially targeted nitroxide radical antioxidants (TEMPOL, MitoTEMPO) were insufficient to ameliorate DU toxicity with or without co-exposure of Sennoside A, suggesting a ROS-independent mechanism.

8. SUPPLEMENTAL FIGURES

**Supplemental Figure 1.**

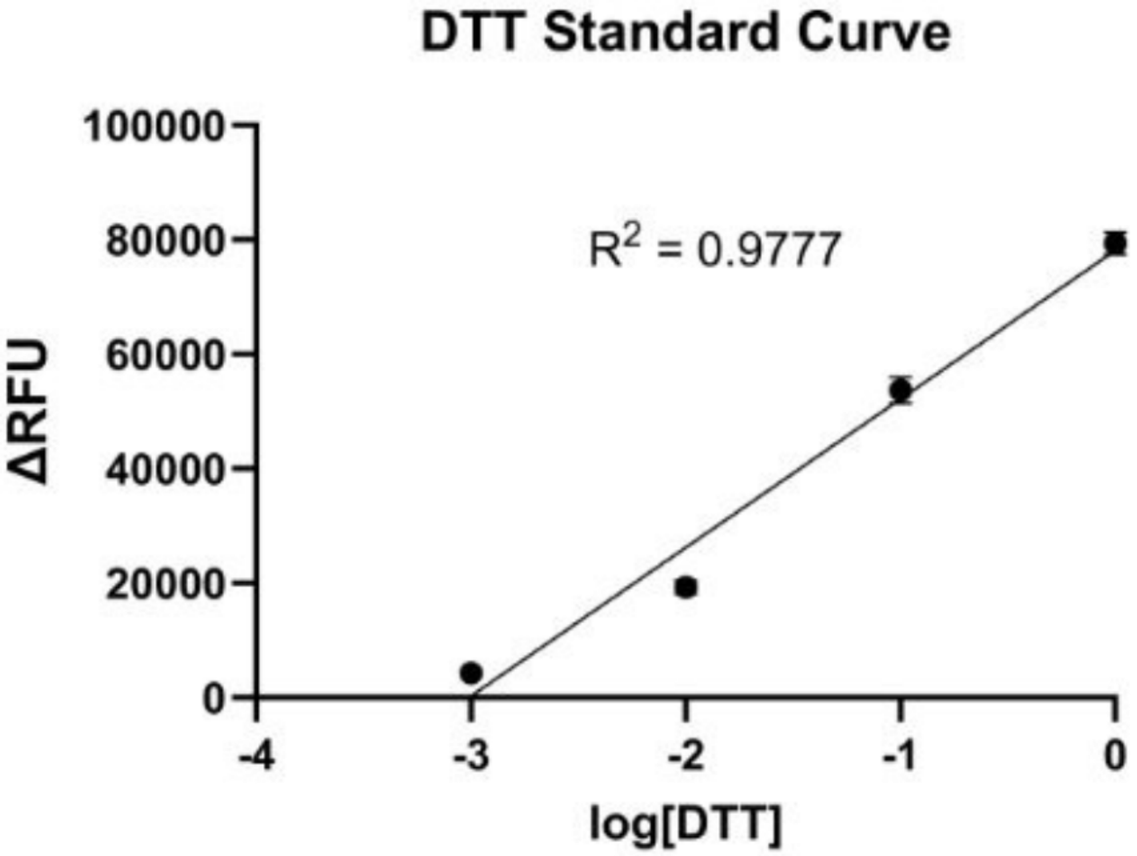

**Supplemental Figure 2.**

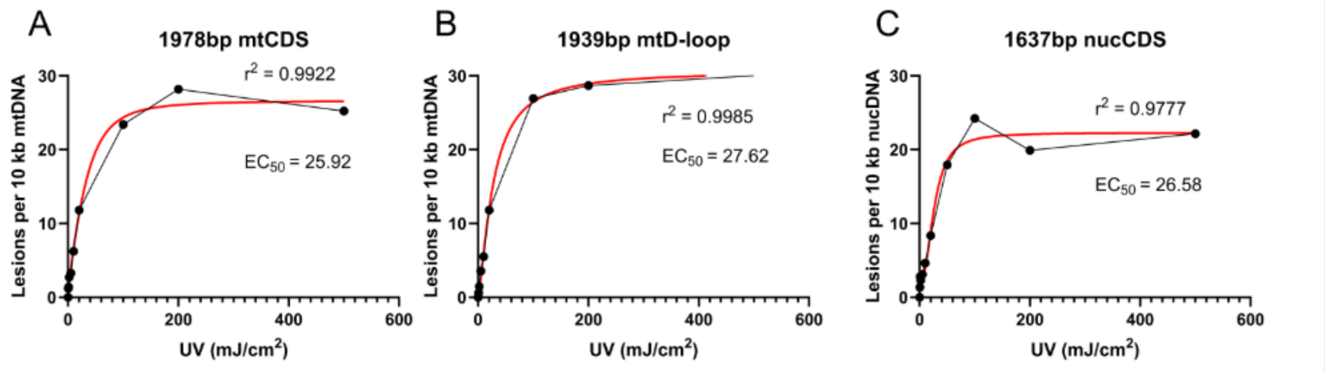


D

| 3669bp mtCDS | 18 ppb | | | | | | 30ppb | | | | | |
|---|---|---|---|---|---|---|---|---|---|---|---|---|
| | p value | trend | mean | std | n-value | strain | p value | trend | mean | std | n-value | strain |
| 1/15/2024 | 0.02239 | | 2.00312 | 0.61934 | 10 | AB | 0.00435 | | 2.03008 | 2.55637 | 13 | AB |
| 9/20/2023 | | | | | 13 | TU | | | | | 9 | TU |
| 8/23/2023 | 0.0061 | down | -0.25838 | 0.38065 | 5 | TU | 0.2329 | none | -0.04825 | 1.42008 | 7 | TU |
| 9/5/2023 | | | | | 7 | TU | | | | | 6 | TU |
| 8/15/2023 | 0.0212 | down | -0.50552 | 0.61541 | 4 | TU | 0.4886 | none | 0.57868 | 0.51561 | 7 | TU |
| 7/3/2023 | 0.0693 | up | 3.40661 | 2.42838 | 5 | TU | 0.0746 | up | 2.95732 | 1.10408 | 7 | TU |
| 6/13/2023 | 0.1341 | none | 5.71847 | 1.34704 | 11 | TU | 0.4359 | none | 6.42874 | 0.96845 | 8 | TU |
| 5/23/2023 | 0.0001 | down | | | | TU | 0.7248 | none | 0.75465 | 1.52288 | 18 | TU |
| 5/15/2023 | 0.412 | none | | | | TU | 0.0449 | down | 1.95902 | 0.19287 | 20 | TU |
| **1978bp mtCDS** | **18 ppb** | | | | | | **30ppb** | | | | | |
| | p value | trend | mean | std | n-value | strain | p value | trend | mean | std | n-value | strain |
| 1/15/2024 | 0.1868 | | 6.47699 | 1.29862 | 10 | AB | 0.18404 | | 5.09127 | 2.76261 | 13 | AB |
| 9/20/2023 | 0.20438 | | 3.18026 | 1.99406 | 13 | TU | 0.14233 | | 3.68903 | 1.60604 | 9 | TU |
| 8/23/2023 | 0.439 | none | 1.85714 | 2.67301 | 5 | TU | 0.0019 | down | -0.70839 | 1.35482 | 7 | TU |
| 9/5/2023 | 0.1175 | none | -0.045 | 0.87513 | 7 | TU | 0.2197 | none | 1.44798 | 0.87168 | 6 | TU |
| 8/15/2023 | 0.0039 | down | -0.76664 | 0.47437 | 4 | TU | 0.3195 | none | 1.17651 | 1.22467 | 7 | TU |
| 7/3/2023 | 0.433 | none | 3.14294 | 3.63338 | 5 | TU | 0.4871 | none | 2.85309 | 2.8801 | 7 | TU |
| 6/13/2023 | 0.3121 | none | 9.86955 | 0.66341 | 11 | TU | 0.1364 | none | 11.209 | 0.63738 | 8 | TU |
| 5/23/2023 | 0.0001 | down | | | | TU | 0.0041 | up | 2.96447 | 2.74837 | 18 | TU |
| 5/15/2023 | 0.3522 | none | | | | TU | 0.064 | up | -3.46868 | 3.20136 | 20 | TU |
| **1939bp mtDLoop** | **18 ppb** | | | | | | **30ppb** | | | | | |
| | p value | trend | mean | std | n-value | strain | p value | trend | mean | std | n-value | strain |
| 1/15/2024 | 0.40519 | | 7.30981 | 2.04005 | 10 | AB | 0.00791 | | 4.29166 | 4.34302 | 13 | AB |
| 9/20/2023 | 0.11127 | | 3.86993 | 2.40458 | 13 | TU | 0.10602 | | 4.96333 | 4.65755 | 9 | TU |
| 8/23/2023 | 0.0226 | up | | | 5 | TU | 0.0444 | up | | | 7 | TU |
| 9/5/2023 | 0.0033 | up | 1.72837 | 0.99082 | 7 | TU | 0.0003 | up | 3.11614 | 1.16716 | 6 | TU |
| 8/15/2023 | 0.0312 | down | -0.40275 | 0.79793 | 4 | TU | 0.4428 | none | 1.529 | 1.56685 | 7 | TU |
| 7/3/2023 | 0.0398 | up | 10.1217 | 8.6412 | 5 | TU | 0.0827 | up | 5.24346 | 2.2736 | 7 | TU |
| 6/13/2023 | 0.1852 | none | 7.5373 | 0.8674 | 11 | TU | 0.2548 | none | 7.87992 | 0.52356 | 8 | TU |
| 5/23/2023 | 0.0479 | down | | | | TU | 0.5393 | none | 7.37746 | 5.84712 | 18 | TU |
| 5/15/2023 | 0.1459 | none | | | | TU | 0.0341 | up | -1.89965 | 3.93444 | 20 | TU |

D (cont.)

| mtDNAcn | 18 ppb | | | | | | 30ppb | | | | | |
|---|---|---|---|---|---|---|---|---|---|---|---|---|
| | p value | trend | mean | std | n-value | strain | p value | trend | mean | std | n-value | strain |
| 1/15/2024 | 0.00561 | down | 797.549 | 368.673 | 10 | AB | 0.00029 | down | 501.559 | 293.699 | 13 | AB |
| 9/20/2023 | 0.47979 | | 452.679 | 195.76 | 13 | TU | 0.20323 | | 355.128 | 141.243 | 9 | TU |
| 8/23/2023 | 0.0443 | down | 269.755 | 57.0121 | 5 | TU | 0.0019 | down | 228.55 | 96.9907 | 7 | TU |
| 9/5/2023 | 0.3051 | none | 341.38 | 54.0545 | 7 | TU | 0.1816 | none | 361.721 | 61.1276 | 6 | TU |
| 8/15/2023 | 0.3432 | none | 206.914 | 92.7806 | 4 | TU | 0.1354 | none | 307.556 | 95.1214 | 7 | TU |
| 7/3/2023 | 0.0019 | up | 393.018 | 58.6769 | 5 | TU | 0.1188 | none | 371.277 | 525.047 | 7 | TU |
| 6/13/2023 | 0.0111 | up | 382.221 | 128.846 | 11 | TU | 0.0127 | up | 410.126 | 180.828 | 8 | TU |
| 5/23/2023 | 0.0051 | down | | | | TU | 0.0148 | down | 289.531 | 219.976 | 18 | TU |
| 5/15/2023 | 0.3248 | none | | | | TU | 0.365 | none | | | 20 | TU |
| **1637bp nucCDS** | **18 ppb** | | | | | | **30ppb** | | | | | |
| | p value | trend | mean | std | n-value | strain | p value | trend | mean | std | n-value | strain |
| 1/15/2024 | 9.6E-06 | up | 5.93303 | 1.9913 | 10 | AB | 2.1E-06 | up | 4.24121 | 0.59427 | 13 | AB |
| 9/20/2023 | 0.35614 | | 3.43919 | 2.23009 | 13 | TU | 0.26868 | | 3.04725 | 2.75495 | 9 | TU |
| 8/23/2023 | 0.3319 | none | 2.20902 | 1.0512 | 5 | TU | 0.1148 | none | 0.58131 | 1.02502 | 7 | TU |
| 9/5/2023 | 0.2358 | none | 1.84871 | 2.08596 | 7 | TU | 0.0674 | down | 0.38924 | 1.40852 | 6 | TU |
| 8/15/2023 | 0.4112 | none | 0.11807 | 2.54076 | 4 | TU | 0.4938 | none | 0.40802 | 1.36991 | 7 | TU |
| 7/3/2023 | 0.3504 | none | 1.8163 | 1.39526 | 5 | TU | 0.0695 | up | 6.70899 | 7.53516 | 7 | TU |
| 6/13/2023 | 0.3946 | none | 10.3406 | 4.90969 | 11 | TU | 0.1162 | none | 11.1576 | 1.78439 | 8 | TU |
| 5/23/2023 | 0.9766 | none | | | | TU | 0.0028 | up | | | 18 | TU |
| 5/15/2023 | 0.3853 | none | | | | TU | 0.163 | none | 0.25073 | 0.19235 | 20 | TU |
| **3502bp nucCDS** | **18 ppb** | | | | | | **30ppb** | | | | | |
| | p value | trend | mean | std | n-value | strain | p value | trend | mean | std | n-value | strain |
| 1/15/2024 | 0.21469 | none | 2.02627 | 1.25959 | 10 | AB | 0.27079 | none | 1.94913 | 1.4946 | 13 | AB |
| 9/20/2023 | 0.09475 | none | 3.07695 | 0.74018 | 13 | TU | 0.16965 | none | 2.7018 | 0.65608 | 9 | TU |
| 8/23/2023 | 0.0036 | up | 4.06733 | 1.89109 | 5 | TU | 0.1291 | none | 0.99083 | 1.07302 | 7 | TU |
| 9/5/2023 | 0.3128 | none | 2.39592 | 0.97626 | 7 | TU | 0.0459 | up | 4.87476 | 2.90437 | 6 | TU |
| 8/15/2023 | 0.1461 | none | -0.66918 | 1.27292 | 4 | TU | 0.0433 | up | -0.54363 | 0.61867 | 7 | TU |
| 7/3/2023 | 0.1209 | none | 4.7562 | 0.68809 | 5 | TU | 0.4983 | none | -6.66528 | 10.0442 | 7 | TU |
| 6/13/2023 | 0.3894 | none | 5.44091 | 1.41391 | 11 | TU | 0.0022 | up | 8.23903 | 1.30417 | 8 | TU |
| 5/23/2023 | 0.08 | up | | | | TU | 0.0001 | up | 6.72149 | 4.07597 | 18 | TU |
| 5/15/2023 | 0.0095 | up | | | | TU | 0.2164 | none | -1.21674 | 3.52797 | 20 | TU |

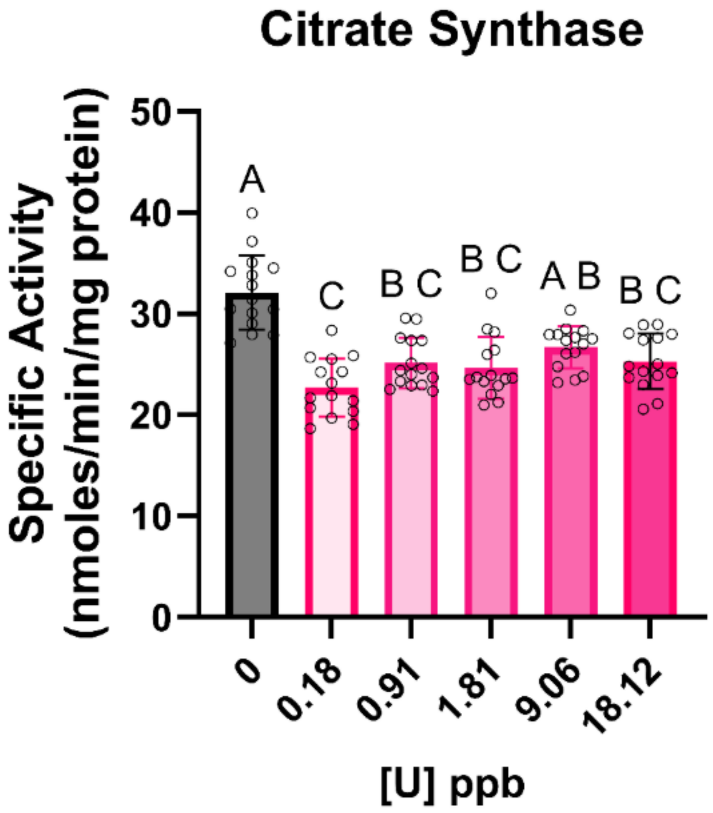


**Supplemental Figure 3.**

**Supplemental Figure 4.**

| dose uranyl nitrate (µg/L) | theoretical $^{238}U$ ppb | measured $^{238}U$ ppb |
|---|---|---|
| 0.3 | 0.1422 | 0.53683 |
| 3 | 1.422 | 1.11683 |
| 30 | 14.22 | 11.22452 |
| 49.7 | 23.5578 | 15.36074 |
| 300 | 142.2 | 26.63082 |
| 3000 | 1422 | 1678.22566 |
| 0.3X Danieau's Solution | 0 | 0.00067 |
| MilliQ-Water | 0 | 0.00044 |
| Blank | 0 | 0.00001 |
| | higher than theoretical | |
| | lower than theoretical | |

9. SUPPLEMENTAL TABLES

**Supplemental Table 1.**

| Gene Target | Direction | Primer Sequence (5' to 3') | Source |
|---|---|---|---|
| *gss* | Forward | GCCAGCAGCACTCTTTCATC | Bian et al. 2023 |
| | Reverse | AACTGTGATTCTGGCTGACCC | |
| *gstp* | Forward | CGACTTGAAAGCCACCTGTGTC | Marques et al. 2024 |
| | Reverse | CTGTCGTTTTTGCCATATGCAGC | |
| *nqo1* | Forward | AGGTGGAGCAGGCGGATCTA | PrimerBlast (intron including) |
| | Reverse | TGGGTACTGCAAACTCATTACAAAG | |
| *ssh1a* | Forward | GGGATGGCACTGACGTGGA | NCBI Blast |
| | Reverse | GTTCTCTCCCTCTCTGTTGGCTTGT | |
| *ssh1b* | Forward | GCCATGGCTCTGGTGACTCT | NCBI Blast |
| | Reverse | CTGTGGAAGATCCCCTGCGT | |
| *ssh2a* | Forward | CACTCCTTGCCACTGACCCA | NCBI Blast |
| | Reverse | AGGGTAAAAGGTGGGCGGAG | |
| *ssh2b* | Forward | CTCCAAGCGCAACATCCAGC | NCBI Blast |
| | Reverse | AGCAGAGGAGCCATTCCCAC | |
| *β-actin* | Forward | CAACAGAGAGAAGATGACACAGATCA | Marques et al. 2024 |
| | Reverse | GTCACACCATCACCAGAGTCCATCAC | |
| *ef1a* | Forward | CTTCTCAGGCTGACTGTGC | Salanga et al. 2019 |
| | Reverse | CCGCTAGCATTACCCTCC | |

**Supplemental Table 2.**

| Primer Name | Sequence (5' to 3') | Amplicon Size (bp) | Amplicon Region | PCR Efficiency (%) | $R^2$ | Melting Temperature (°C) | Position on the mt Genome (bp) |
|---|---|---|---|---|---|---|---|
| mtFp 3 | TTAAAGCCCCGAATCCAGGTGAGC | 3669 | coding region mtDNA | 87.7 | 0.981 | 85.0 | 2303 - 5971 |
| mtRp 3 | TTAGGGGTAGTGAGTTTTGGGTC | | | | | | |
| mtFp 4 | GGATTCCAAGACGCAGCATCACCTG | 1978 | coding region mtDNA | 97.1 | 0.997 | 84.5 | 8151 - 10128 |
| mtRp 4 | GGAGCGGCACTTCAAATGGGTCAAG | | | | | | |
| mtFp 5 | CCTTACACGATTCTTCGCATTCCAC | 1939 | D-loop mtDNA | 89 | 0.999 | 83.0 | 15829 - 1171 |
| mtRp 5 | GGCTTGGCTAGGCGTCTTGG | | | | | | |
| mtFp 6 | CGAGGAGCAGGTATCAGGCACA | 55 | coding region mtDNA | 88.6 | 0.997 | 81.0 | 1119 - 1173 |
| mtRp 6 | GTGGCTTGGCTAGGCGTCTTG | | | | | | |
| nucFp 7 | GTCCTTGCAGGTTGGCAAATGG | 3502 | AHR1 gene, nucDNA | 104 | 0.995 | 88.0 | n/a |
| nucRp 7 | GACCTTGTCTGGTTTTCATCCC | | | | | | |
| nucFp 8 | TCATCCTGTTATCCACCACACTGTTG | 1637 | AHR1 gene, nucDNA | 82.2 | 0.985 | 85.0 | n/a |
| nucRp 8 | TGGTTCTTGGCTACACTGAGATTGAG | | | | | | |
| nucFp 9 | CCAAGGTCCGACATAACTCACTTCTG | 55 | AHR1 gene, nucDNA | 104 | 0.997 | 78.5 | n/a |
| nucRp 9 | GACATGATGTACTGTGCTGACAACCA | | | | | | |
